# Copy number variants underlie the major selective sweeps in insecticide resistance genes in *Anopheles arabiensis* from Tanzania

**DOI:** 10.1101/2024.03.11.583874

**Authors:** Eric R. Lucas, Sanjay C. Nagi, Bilali Kabula, Bernard Batengana, William Kisinza, Alexander Egyir-Yawson, John Essandoh, Sam Dadzie, Joseph Chabi, Arjen E. Van’t Hof, Emily J. Rippon, Dimitra Pipini, Nicholas J. Harding, Naomi A. Dyer, Chris S. Clarkson, Alistair Miles, David Weetman, Martin J. Donnelly

**Affiliations:** Department of Vector Biology, Liverpool School of Tropical Medicine, Pembroke Place, Liverpool L3 5QA, UK; National Institute for Medical Research, Amani Research Centre, P.O. Box 81, Muheza, Tanzania; Department of Biomedical Sciences, University of Cape Coast, Cape Coast, Ghana; Department of Parasitology, Noguchi Memorial Institute for Medical Research, University of Ghana, Accra, Ghana; Biology Centre of the Czech Academy of Sciences, Institute of Entomology, Branišovská 31, 370 05 České Budějovice, Czech Republic; Big Data Institute, Li Ka Shing Centre for Health Information and Discovery, University of Oxford, Oxford, United Kingdom; Wellcome Sanger Institute, Hinxton, Cambridge CB10 1SA, United Kingdom

**Keywords:** Carboxylesterase, GWAS, insecticide resistance, genomic surveillance, *Anopheles*

## Abstract

To keep ahead of the evolution of resistance to insecticides in mosquitoes, national malaria control programmes must make use of a range of insecticides, both old and new, while monitoring resistance mechanisms. Knowledge of the mechanisms of resistance remains limited in *Anopheles arabiensis*, which in many parts of Africa is of increasing importance because it is apparently less susceptible to many indoor control interventions. Furthermore, comparatively little is known in general about resistance to non-pyrethroid insecticides such as pirimiphos-methyl (PM), which are crucial for effective control in the context of resistance to pyrethroids. We performed a genome-wide association study to determine the molecular mechanisms of resistance to deltamethrin (commonly used in bednets) and PM, in *An. arabiensis* from two regions in Tanzania. Genomic regions of positive selection in these populations were largely driven by copy number variants (CNVs) in gene families involved in resistance to these two insecticides. We found evidence of a new gene cluster involved in resistance to PM, identifying a strong selective sweep tied to a CNV in the *Coeae2g-Coeae6g* cluster of carboxylesterase genes. Using complementary data from *An. coluzzii* in Ghana, we show that copy number at this locus is significantly associated with PM resistance. Similarly, for deltamethrin, resistance was strongly associated with a novel CNV allele in the *Cyp6aa* / *Cyp6p* cluster. Against this background of metabolic resistance, target site resistance was very rare or absent for both insecticides. Mutations in the pyrethroid target site *Vgsc* were at very low frequency in Tanzania, yet combining these samples with three *An. arabiensis* individuals from West Africa revealed a startling diversity of evolutionary origins of target site resistance, with up to 5 independent origins of *Vgsc*-995 mutations found within just 8 haplotypes. Thus, despite having been first recorded over 10 years ago, *Vgsc* resistance mutations in Tanzanian *An. arabiensis* have remained at stable low frequencies. Overall, our results provide a new copy number marker for monitoring resistance to PM in malaria mosquitoes, and reveal the complex picture of resistance patterns in *An. arabiensis*.

## 1. Introduction

The evolution of insecticide resistance in disease vectors threatens effective control of vector-borne diseases such as malaria (Hemingway et al., 2016; Kafy et al., 2017; Protopopoff et al., 2018; Maiteki-Sebuguzi et al., 2023), in the same way as antibiotic resistance is jeopardising the effective treatment of bacterial infections. In large parts of Africa, malaria-transmitting mosquitoes have already developed resistance to the most widely-used class of public health insecticides, pyrethroids (Hancock et al., 2020, 2022). In response to this, other insecticides have been deployed, such as indoor residual spraying (IRS) with the organophosphate pirimiphos-methyl (PM) (Abong’o et al., 2020). For the effectiveness of these interventions to be sustained, resistance to the new compounds needs to be anticipated and monitored. High levels of PM resistance have already been detected in parts of West Africa, where it is primarily driven by a single nucleotide polymorphism (SNP) and copy number variation (CNV) in the target site of PM, *Ace1* (Grau-Bové et al., 2021). In contrast, researchers in East Africa have reported fewer cases of PM resistance (Supplementary Fig. S1), and an absence of *Ace-1* resistance mutations in any malaria vector species. It is therefore crucial to investigate populations showing early evidence of PM resistance to understand the nature of this resistance and unravel the genetic mechanisms that underlie it, to better monitor incipient resistance across the region.

While indoor-based interventions such as IRS and insecticide-treated nets (ITNs) have successfully reduced numbers of the major vector *Anopheles gambiae s*.*s*. in East Africa, outdoor-biting species such as *An. arabiensis* have been less affected (Okumu and Finda, 2021). Resistance levels in *An. arabiensis* are typically lower than in indoor biting species, probably due to their reduced exposure to insecticides, but is nonetheless appreciable to some active ingredients (Pinda et al., 2020; Orondo et al., 2021; Matiya et al., 2022; Mawejje et al., 2023). This is a cause for concern since *An. arabiensis* is a significant vector of malaria (Okumu and Finda, 2021), and in some areas the primary vector (Matowo et al., 2014; Degefa et al., 2021; Mwalimu et al., 2024). In East Africa, resistance in *An. arabiensis* has been reported to deltamethrin (Pinda et al., 2020; Matiya et al., 2022), a pyrethroid widely used in bednets, and PM (Omoke et al., 2023). This provides an ideal opportunity to study the genomics of resistance in this species, both for established (deltamethrin) and recently introduced (PM) insecticides.

As part of the Genomics for African *Anopheles* Resistance Diagnostics (GAARD) project, we are using large scale whole genome sequencing to investigate the genomics of insecticide resistance in key regions of Africa. Here we investigated resistance in *An. arabiensis* from two contrasting regions of Tanzania (Fig. 1a). Moshi is an elevated area with extensive rice and sugarcane plantation and associated irrigation (Ijumba, Mosha and Lindsay, 2002), with the possibility of resistance developing due to exposure to insecticides used on crops. Muleba is an area that has been the site of vector control trials, and where mosquitoes have thus been exposed to a range of public health insecticides, including PM (West et al., 2014; Kisinza et al., 2017; Protopopoff et al., 2018). We conducted a genome-wide association study (GWAS) of resistance to deltamethrin and PM in these two populations.

**Figure 1:**
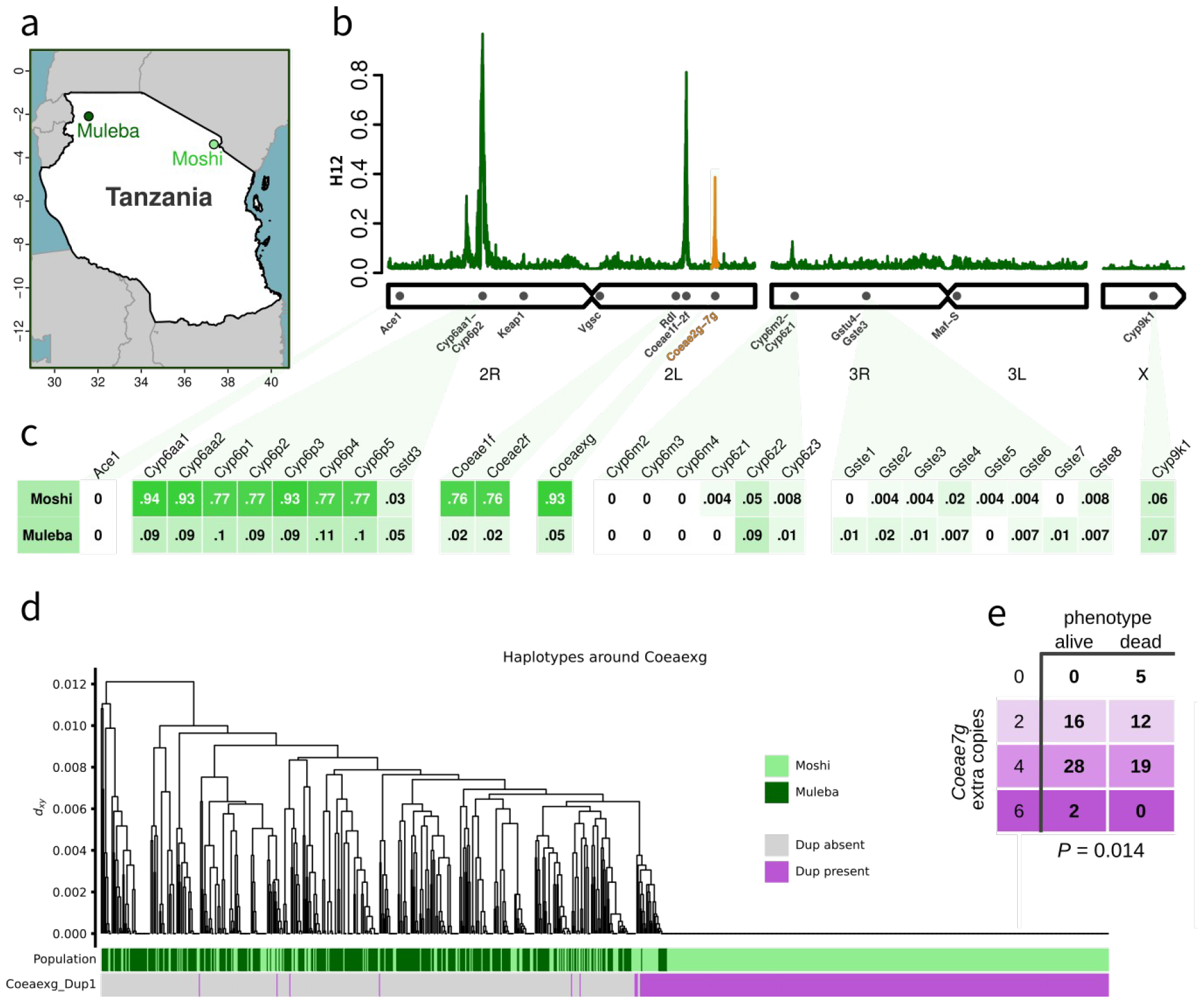
**A**. Map of sampling locations. GPS coordinates are given in Supplementary Data S1. **B**. Genome-wide H12 calculated in 2000 SNP windows in samples from Moshi, showing peaks in selection signal around Cyp6aa/Cyp6p, Coeae1f-2f and Coeae2g-7g (highlighted in orange). **C**. Proportion of samples carrying a CNV in key metabolic genes (columns) in each of our two sample sites (rows). Mosquitoes in Moshi had > 70% frequency of CNVs in the Coeae1f-2f and Coeae2g-6g genes (averaged as Coaeaxg in the table), as well as Cyp6aa/Cyp6p. **D**. Haplotype clustering of the genomic region around Coeaexg in Moshi and Muleba. Haplotypes bearing the CNV allele Coeaexg_Dup1 were almost perfectly associated with the large swept cluster seen on the right, indicating that the CNV is likely to be driving the selective sweep. **E**. In An. coluzzii from Korle-Bu, Ghana, copy number of Coeae7g was significantly associated with resistance to PM (P = 0.014 after controlling for CNV in Ace1).

## 2. Results

### 2.1. Bioassays

Preliminary bioassays conducted in Moshi and Muleba indicated the presence of deltamethrin resistance in both locations, but PM resistance only in Moshi (Supplementary Data S1). In Moshi, 24hr mortality to deltamethrin ranged from 53% at 0.5x the WHO diagnostic concentration, to 73% at 2.5x, then 99-100% at each of 5x, 7.5x and 10x. Mortality was slightly higher in Muleba, with 64% mortality at 0.5x, 71% at 1x, 97% at 2.5x, then 100% at 5x and above. For PM, in Moshi mortality ranged from 58% at 0.5x to 86% at 1x, then 100% at 2x. In contrast, there was no evidence of any resistance to PM in Muleba, with 100% mortality even at 0.5x concentration. These bioassays were conducted on mosquitoes identified morphologically as *An. gambiae s*.*l*.. Molecular species identification performed on a subset of these confirmed that all samples in Muleba (196 out of 196) and nearly all in Moshi (382 out of 384) were *An. arabiensis*, with the final two in Moshi being *An. gambiae s*.*s*. All further analyses were performed on *An. arabiensis* only.

### 2.2. Overview of genomic data

Samples for this study were sequenced as part of the MalariaGEN Vector Observatory release Ag3.7 (https://www.malariagen.net/data). Data were obtained from 467 individual female mosquitoes across 3 sample sets (Table 1). The phenotype of each individual was defined by whether they were alive after exposure to a high dose of insecticide (“resistant”) or dead after exposure to a lower dose (“susceptible”), thus providing strong phenotypic separation between phenotypes. Exposure conditions to generate the distinct phenotype classes were calculated separately for each location, and are reported in Supplementary Data S1.

**Table 1:**
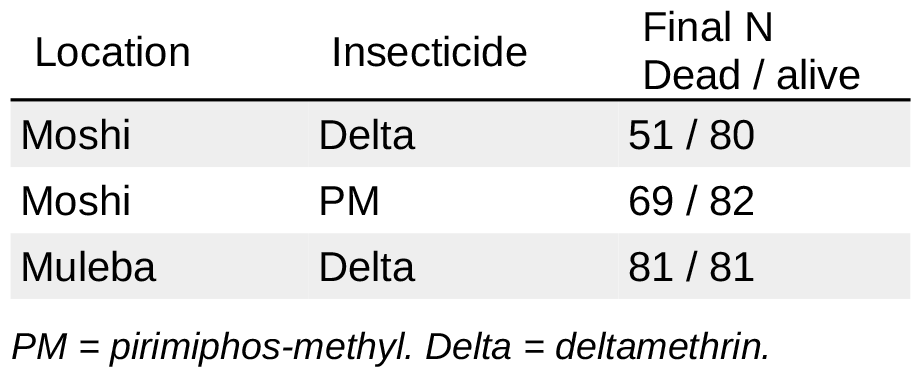
Number of samples sequenced in each of the three sample sets (rows), after removal of siblings and contaminated samples.

We calculated kinship using the KING statistic (Manichaikul et al., 2010) pairwise across all 467 samples to identify close kin pairs (full sibs), which would be non-independent data points in an association study. This resulted in the identification of 18 sib groups containing a total of 38 individuals (16 groups of two siblings, 2 groups of three siblings). All sib groups contained only samples from a single location (2 groups from Moshi, 16 groups from Muleba). Depending on the analysis (see methods), we either discarded all but one randomly chosen individual per sib group per sample set (thus removing 19 samples) or performed permutations in which we varied which individuals were discarded in each sib group. We found that four of the samples had universally high relatedness values to all other samples in the dataset. Closer inspection revealed that these samples had elevated heterozygosity caused by cross-sample contamination (Supplementary Fig. S2), leading to inflated KING values. We therefore removed these four samples from all analyses.

A PCA of the samples based on SNP genotypes indicated genetic differentiation between our two sampling sites, but no other evidence of defined clusters in the first 4 principal coordinates (Supplementary Fig. S3).

### 2.3. Signals of selection point to a new carboxylesterase gene cluster

We first identified regions of the genome undergoing recent positive selection by performing genome-wide H_12_ scans (Fig. 1b, Supplementary Fig. S4), combining the data from the deltamethrin and PM experiments. Signals of selection in *Anopheles gambiae* are often the result of insecticidal pressures (*Anopheles gambiae* 1000 Genomes Consortium et al., 2017), but do not indicate which insecticides are responsible for a given signal, and thus constitute a preliminary analysis of the data to identify regions of potential interest.

In both Moshi and Muleba, the strongest signal of selection across the genome is centred on the cluster of *Cyp6aa* / *Cyp6p* genes on chromosome 2R, a region repeatedly associated with resistance to deltamethrin. H_1x_ analysis (Miles, 2021) indicated that the signals in Moshi and Muleba in this genomic region are shared, with the same haplotype(s) underlying the sweep in both regions (Supplementary Fig. S4).

In Moshi, two peaks in H_12_ were also found on chromosome 2L (Fig. 1b). The first was centred on the carboxylesterases *Coeae1f* (AGAP006227) and *Coeae2f* (AGAP006228), which have been implicated in resistance to PM in West Africa (Nagi et al., 2024). The second peak was centred on another carboxylesterase cluster, *Coeae2g* (AGAP006723) - *Coeae6g* (AGAP006727), which has not previously been associated with resistance.

### 2.4. Copy number variants are associated with resistance to PM and deltamethrin

One of the most commonly described methods of insecticide resistance is metabolic resistance (Liu, 2015), where increased levels, activity, or improved affinity, of metabolic enzymes accelerates the detoxification of insecticides and their by-products. Metabolic resistance can be achieved through a much broader range of mutations than target site resistance, making it harder to identify causative alleles. However, a tractable group of mutations that have repeatedly been associated with resistance are Copy Number Variants (CNVs, (Weetman, Djogbenou and Lucas, 2018), which increase the number of genomic copies of a gene.

CNVs in the carboxylesterase genes *Coeae1f* and *Coeae2f* (which we collectively refer to as *Coeaexf*) have recently been associated with resistance to PM in *An. gambiae s*.*s*. from Ghana, and have been found in *An. arabiensis* from Tanzania (Nagi et al., 2024). In our samples, we found the previously identified *Coeaexf*_Dup2 CNV allele in 74% of samples from Moshi, and 1% of samples from Muleba (Fig. 1B, Supplementary Fig. S5). We also found 5 other CNV alleles in this cluster, all at low frequencies ranging from 0.4% to 4% of samples in either Moshi or Muleba (Supplementary Fig. S5). Haplotype clustering of the *Coeaexf* region indicated the presence of two selective sweeps, with the more common of the two sweeps being associated with *Coeaexf*_Dup2 (Supplementary Fig. S6). However, *Coeaexf*_Dup2 was present in only a subset of the haplotypes in the sweep, indicating that this CNV likely appeared on this haplotype after it began sweeping. There was no association of *Coeaexf* copy number in Moshi with resistance to either deltamethrin (*P* = 0.96 and *P* = 0.83 for *Coeae1f* and *Coeae2f* respectively) or PM (*P* = 0.28 and *P =* 0.29). Because the lack of association with PM resistance was unexpected, given the role that *Coeaexf* CNVs play in PM resistance in Ghana, we investigated whether this could be due to lack of statistical power in our data, but this was not the case. We ran simulations assuming that presence / absence of the CNV provided a similar effect size of resistance as was previously found in *An. gambiae* from Ghana (Nagi et al., 2024) and found that we had 88% power to detect the effect in our data.

To explore the selection signal which we identified in the carboxylesterase genes *Coeae2g*-*Coeae6g* (which we will refer to as *Coeaexg*), we also investigated CNVs in this genetic region (Fig. 1c, Supplementary Fig. S5). We found a CNV, which we call *Coeaexg*_Dup1 (Supplementary Fig. S7), at very high frequency (94% of samples) in Moshi (where resistance to PM is prevalent) and lower frequency (6%) in Muleba (where mosquitoes are completely susceptible to PM). Haplotype clustering indicated the presence of one major swept haplotype in this genomic region, which corresponded almost exactly to the presence of the CNV (Fig. 1d), implying that the CNV is driving the selective sweep. Copy number of this CNV was highly variable and could reach very high values (the median copy number among samples carrying the CNV was 8, with a maximum of 28 extra copies). However, we found no significant association of copy number in Moshi with resistance to either deltamethrin (*P* = 0.94) or PM (*P* = 0.38).

In a previous GAARD project study in West Africa, we had identified a signal of association with PM resistance in *An. coluzzii* on chromosome 2L in the regions of 36898300-37190282 and 37558030-37585789 (Lucas et al., 2023). These regions did not include the *Coeaexg* gene cluster itself (2L:37282290-37295276) and the signals had not been prioritised for further investigation. In light of the current observation, we revisited the West African GAARD data and searched for CNVs in *Coeaexg*. We found CNVs at low frequency in *An. gambiae* populations from Madina and Obuasi (Ghana) and Baguida (Togo), as well as in *An. coluzzii* from Avrankou (Benin). In contrast, in *An. coluzzii* from Ghana (Korle-Bu), we found high CNV frequencies comparable to those in Moshi (94% of samples), although the copy number of these CNVs was lower than in Moshi (median: 4, max: 6 extra copies). The CNVs in West Africa were less clearly defined, in terms of discordant reads that could precisely distinguish CNV alleles and identify start and end points, but they encompassed a larger genomic region than the *An. arabiensis* allele, including *Coeae7g* (Supplementary Fig. S7). There was a significant association of *Coeaexg* CNVs with resistance to PM in Korle-Bu. Copy number of all carboxylesterases within the CNV (*Coeae2g - Coeae7g*) were highly cross-correlated, and thus when the copy number of one gene was included, the addition of other genes did not further improve the model. The *Coeaexg* gene with the strongest association was *Coeae7g*, both with the marker alone in the model (*P* = 0.031) and after inclusion of copy number in *Ace1* (*Ace1 P* = 3×10^−10^; *Coaea7g P* = 0.014, Fig. 1e). However, we note that all the genes in the cluster were significantly associated with PM resistance in a model containing only that gene and *Ace1* (eg: *Coeae6g, P* = 0.02). From these data alone, it is therefore uncertain which proteins in this cluster are most important for conferring resistance.

CNVs in the *Cyp6aa / Cyp6p* region were at much higher frequency in Moshi (77 - 94% depending on the gene) compared to Muleba (9 - 11%, Fig. 1B). Only four samples, all from Moshi, carried one of the 30 CNV alleles previously identified in phase 3 of the Ag1000G project (Supplementary Fig. S5). The remaining CNVs comprised 7 new alleles which we named *Cyp6aap*_Dup31 - *Cyp6aap*_Dup37). The most common alleles were Dup33 (found in 77% of Moshi samples and 7% of Muleba samples) and a pair of CNVs in complete linkage with each-other, Dup31 and Dup32 (33% of Moshi samples). We investigated whether the haplotype undergoing a selective sweep in this genomic region was associated with these CNVs. A haplotype clustering tree of the region showed a large selective sweep, shared between Moshi and Muleba (cluster 1 in Supplementary Fig. S6). Both the Dup31/32 and Dup33 CNVs formed separate sub-groups within this haplotype cluster, indicating that they likely appeared on this haplotype after it began sweeping. A second, smaller, selective sweep was also seen (cluster 2 in Supplementary Fig. S6). Few haplotypes belonged to neither sweep, indicating that haplotypes in *Cyp6aa* / *Cyp6p* that have been under positive selection are now almost fixed in the population.

Copy number of genes in the *Cy6paa/Cyp6p* region was significantly associated with resistance to deltamethrin in Muleba, but not in Moshi. As with *Coeaexg*, all *Cyp6aa/Cyp6p* genes were highly correlated with each-other in terms of copy number, and thus it is impossible to confidently determine from these data which gene was of primary importance. Generalised linear models of phenotype association found that *Cyp6p2* and *Cyp6p3* showed the strongest association of copy number with resistance (*P* = 8×10^−6^, compared to *P* = 2×10^−4^ for *Cyp6aa1*), and after inclusion of one of these genes in the model, no other genes provided further significant improvement. We then investigated the two CNV alleles (*Cyp6aap*_Dup31/32 and *Cyp6aap*_Dup33) separately, as well as the swept haplotype. As with copy number, *Cyp6aap*_Dup33 was strongly associated with resistance in Muleba (*P* = 3×10^−5^, *Cyp6aap*_Dup31/32 was absent in Muleba), but not Moshi (*P* = 0.1 for *Cyp6aap_Dup33, P* = 0.48 for *Cyp6aap*_Dup31/32). In both locations, the large swept haplotype itself was nearly, but not quite, significantly associated with resistance (*P* = 0.084 and *P* = 0.08 in Moshi and Muleba respectively), suggesting that in Muleba, *Cyp6aap*_Dup33 provides resistance over and above that which might be conferred by other mutations on its haplotype background.

### 2.5. Known resistance SNPs

Target site resistance mutations were very rare in our samples. In *Ace1* (the target site of PM), the only known resistance SNP in *An. gambiae s*.*l*. is *Ace1*-280S, and this was completely absent. We found five non-synonymous SNPs in *Ace1* with a minor allele count of at least 5 in our PM sample set from Moshi, but all were low frequency and none were significantly associated with resistance (Supplementary Table S1).

The only recognised resistance SNP present in *Vgsc* (target site of deltamethrin) was *Vgsc*-995F (2 out of 564 haplotypes in Moshi, 0 out of 324 haplotypes in Muleba). We also found the SNP *Vgsc*-1507I, which has previously been found on the haplotype background of *Vgsc*-995F in *An. coluzzii* from Guinea (Clarkson et al., 2021), in a single sample from Moshi, which did not carry *Vgsc*-995F. The most common target site resistance mutation was in *Rdl*, the target site for the organochlorine dieldrin, with *Rdl*-296S found in 2% of haplotypes from Moshi. In contrast to these target site mutations, the metabolic resistance SNP *Cyp4j5*-43F was fixed for the mutant allele in all our samples.

We further investigated the two *Vgsc*-995F mutants to determine whether they were introgressed from, or of similar evolutionary origin to, the same mutation found in other populations of *An. gambiae* and *An. arabiensis*. We performed haplotype clustering in the *Vgsc* gene as in previous work (*Anopheles gambiae* 1000 Genomes Consortium et al., 2017; Clarkson et al., 2021), combining our samples with all 2784 samples from phase 3 of the Ag1000G. This includes 368 *An. arabiensis* individuals, but only three of which (*i*.*e*., six haplotypes) are from West Africa. All three are from Burkina Faso and have resistance mutations in *Vgsc* (2 *Vgsc*-995F and 4 *Vgsc*-995S). We found little geographical structure among *Vgsc* haplotypes, with the six haplotypes from Burkina Faso being interspersed among East African samples (Supplementary Fig. S8). Despite there being only four *Vgsc*-995F haplotypes in the entire sample set, these were found in three different parts of the haplotype tree, indicating that between them they represent three different evolutionary origins of the mutation, with the two mutant haplotypes in our Tanzanian samples being of different origins, and the two Burkinabè mutants together forming a third origin. The four *Vgsc*-995S haplotypes from Burkina Faso formed two clusters which, while being close together in the tree, were separated by wild-type haplotypes. None of the *Vgsc*-995 mutations in *An. arabiensis* clustered with *An. gambiae* s.s. or *An. coluzzii* haplotypes, indicating that these are likely to have originated within *An. arabiensis* rather than having introgressed from these other species.

### 2.6. Windowed measures of differentiation / selection to identify genomic regions associated with resistance

We performed agnostic genomic scans of phenotype association as described for a previous analysis (Lucas et al., 2023). This uses F_ST_, PBS, and ΔH_12_ (difference in H_12_ signal between resistant and susceptible subsets) calculated in 1000 SNP windows, as well as identifying 100,000 bp windows with a high frequency of low *P*-value SNPs identified with a SNP-wise GWAS (Fig. 2, Supplementary Data S3-6, Supplementary Fig. S9).

**Figure 2:**
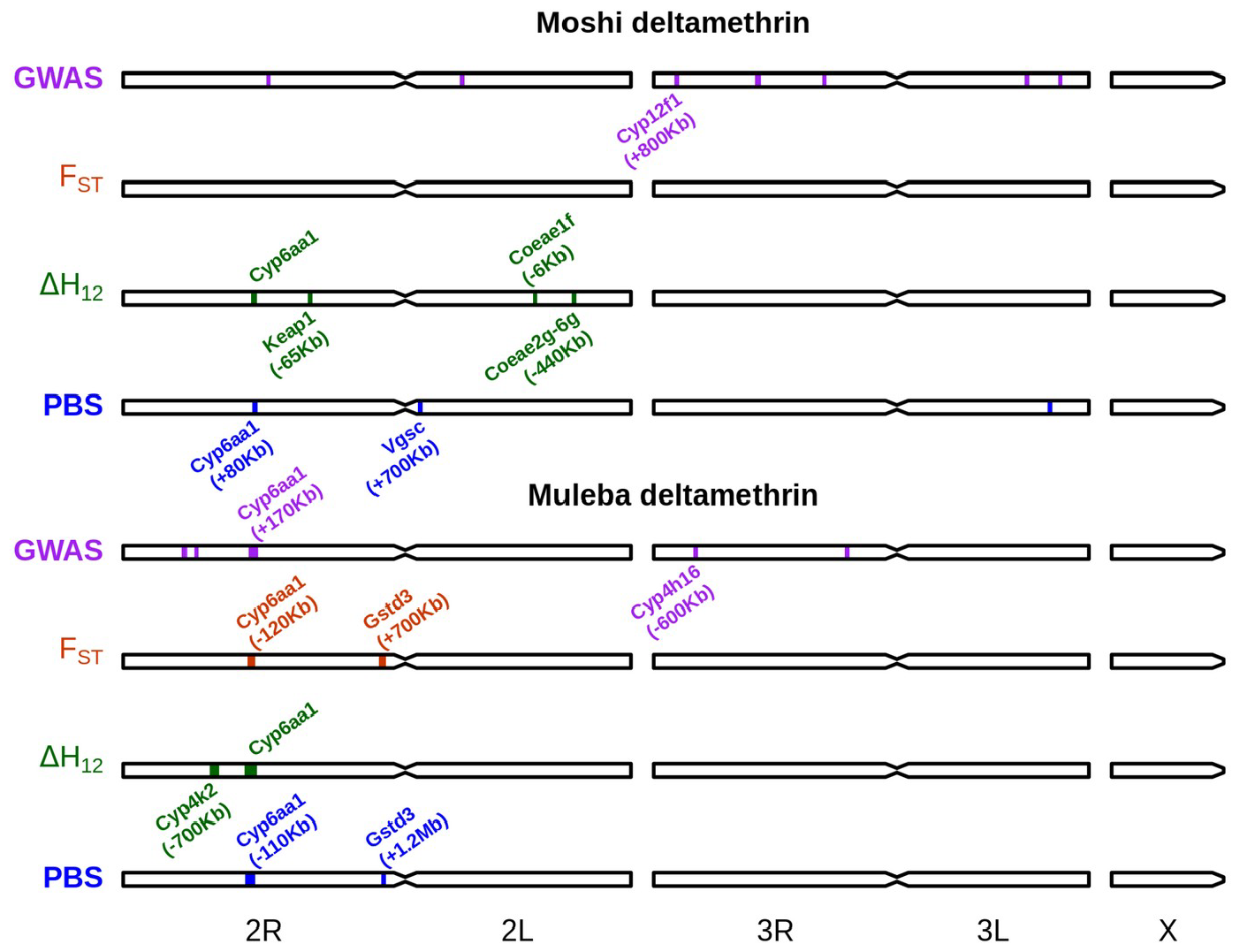
Genomic regions implicated in deltamethrin resistance according to our four approaches (windowed GWAS, FST, ΔH12, PBS). Regions are annotated with genes discussed in the manuscript as possibly causing the signal. Genomic distances in brackets indicate the distance of the peak either to the left (-) or right (+) of the gene in question.

For deltamethrin (Fig. 2), the region around *Cyp6aa1* (*Cyp6aa*/*Cyp6p* region) was consistently associated with resistance in both Moshi and Muleba, and across most methods, although the signal was not always centred directly on this gene cluster, sometimes being as far as 170 Kbp away. Outside of this gene region, two signals of association were near other clusters of cytochrome P450s (*Cyp12f1-4* in Moshi, *Cyp4h16-18* in Muleba) and in Moshi the PBS analysis suggested a region 700 Kbp away from the target site *Vgsc* was associated with resistance. There were ΔH_12_ signals of association in Moshi near *Keap1* (AGAP003645, 65Kbp away) and with both *Coeaexf* (6 Kbp away) and *Coaeaexg* (440 Kbp away). In Muleba, both PBS and F_ST_ detected regions of association with resistance near the end of chromosome 2R, which were respectively 1.2 Mbp and 700 Kbp away from the *Gstd* cluster of glutathione esterase genes. Previous work had demonstrated the presence of CNVs in *Gstd3* in *An. arabiensis* (Tomlinson, 2021) and *Cyp12f1* in *An. gambiae* (Lucas et al., 2019). We therefore investigated whether copy number in these two genes was associated with resistance. We found elevated copy number of *Gstd3* in 5% and 3% of samples from Muleba and Moshi respectively, and copy number was nearly significantly associated with resistance to deltamethrin in Muleba (*P* = 0.052) but not Moshi (*P* = 0.39). When combining both locations together and including location as a random effect, copy number in *Gstd3* reached marginal significance (*P* = 0.046) when it was the only fixed effect term in the model, but this significance disappeared when copy number of *Cyp6aa1* (*P* = 0.0016) was also included, leaving the association of *Gstd3* uncertain. We found no CNVs in *Cyp12f1* in *An. arabiensis*. Revisiting our data from West Africa as above, we did find CNVs in *Cyp12f1* in all populations, but these were not associated with resistance to either deltamethrin or PM.

For PM, there were few windows associated with resistance, and those that were, were not close to any gene families typically associated with resistance. Interestingly, we found a window with a high frequency of low *P*-value SNPs in the region around the *Ace1* gene (340 Kbp away), despite the lack of known resistance SNPs or CNVs in *Ace1* in this population.

## 3. Discussion

We have identified a new cluster of carboxylesterase genes associated with resistance to PM, and possibly deltamethrin, in wild-caught *Anopheles* mosquitoes. A CNV encompassing the genes *Coeae2g*-*Coeae6g* was found at much higher prevalence in Moshi, where PM resistance was prevalent, compared to Muleba, where resistance was absent. Furthermore, a larger CNV in the same gene cluster in *An. coluzzii* from Ghana was significantly associated with survival to PM exposure. Carboxylesterases are a classic example of insecticide resistance driven by copy number variants, with the genes *Est2* and *Est3* in *Culex* (Raymond et al., 2001), and *CCEae3a and CCEae6a* in *Aedes* (Grigoraki et al., 2015; Cattel et al., 2021), showing highly elevated copy number associated with resistance to organophosphates. We similarly found very high copy number of *Coeae2g*-*Coeae6g* in Tanzania, with as many as 26 extra copies of these genes in a single individual, yet curiously there was no significant association of copy number with PM resistance in these samples. One possibility is that the very high frequency of the CNV (being found in 93% of samples in Moshi), led to low statistical power, but we note that copy number was highly variable, ranging from 1 extra copy to 26, and we would therefore expect that this variability in copy number would be associated with resistance and provide sufficient power.

While there was no statistical association of *Coeaexg* and *Coeaexf* copy number with resistance to PM in either of our Tanzanian sites, our agnostic genome-wide scans found evidence of association with deltamethrin resistance near both these gene clusters in Muleba. This is unexpected, as carboxylesterases are not typically associated with resistance to pyrethroids (Poulton et al., 2023), and any evidence of association so far has been correlative (Ishak et al., 2017; Sandeu et al., 2020). Furthermore, in our CNV analysis, copy number of neither gene group was associated with resistance to deltamethrin. Given that the association signals were 6 Kbp away from *Coeaexf*, and 440 Kbp away from *Coeaexg*, it may be that these results are false positives. However, we consider that the presence of two independent signals in related carboxylesterases makes the possibility of false positives unlikely, and this gene cluster is therefore of concern as a potential target of cross-resistance.

As has been found in *An. gambiae* and *An. coluzzii* (Njoroge et al., 2022; Kouamé et al., 2023; Lucas et al., 2023), resistance to deltamethrin in *An. arabiensis* seems to be primarily driven by the *Cyp6aa/Cyp6p* cluster, with this being a consistent conclusion throughout our analysis, from selection scans, GWAS and CNV association studies. In *An. gambiae* and *An. coluzzii*, this metabolic resistance occurs in a context in which target site resistance is largely fixed. In contrast, in *An. arabiensis*, it seems to be the dominant form of resistance, a situation similar to that found in *An. funestus*, where *P450*-based resistance is widespread in the absence of target site mechanisms (Irving and Wondji, 2017; Ibrahim et al., 2018; Weedall et al., 2019; Wamba et al., 2021). The CNV alleles found in *An. arabiensis* are distinct from those in *An. gambiae* and *An. coluzzii*, but similarly provide resistance to deltamethrin. The emergent picture from the *An. gambiae* species complex is thus that metabolic resistance to deltamethrin is consistently driven by mutations in the *Cyp6aa*/*Cyp6p* cluster (Boonsuepsakul, Luepromchai and Rongnoparut, 2008; Ibrahim et al., 2018; Lucas et al., 2023), and that these mutations are very often CNVs in *Cyp6aa1*. These CNVs are however frequently accompanied by other mutations. For example, in our study, both of the CNVs that we found appear on the background of a haplotype undergoing a hard selective sweep, yet only the CNVs, not the haplotype, were significantly associated with deltamethrin resistance in our data, suggesting that the CNVs provide a substantial boost to resistance. In *An. gambiae* from Uganda, Kenya, Tanzania and the Democratic Republic of Congo, a CNV covering only *Cyp6aa1* (*Cyp6aa*_Dup1), again associated with deltamethrin resistance, has spread to near fixation over the course of around 10 years (Njoroge et al., 2022). In a striking parallel with our study, this CNV occurs on the background of a swept haplotype, although the non-CNV version of this haplotype is now so rare that phenotypic analysis of the CNV in isolation from other mutations on the haplotype cannot be performed.

In Ag1000g, a total of 38 CNVs have now been described at the *Cyp6aa/Cyp6p* locus, although many are rare or have not yet been tested for resistance association (The *Anopheles gambiae* 1000 Genomes Consortium, 2021). Over and above this huge diversity of CNVs, other non-CNV mutations are either confirmed or suspected to bring about resistance. In Ghana, a swept haplotype in the *Cyp6aa*/*Cyp6p* cluster was associated with resistance to deltamethrin (Lucas et al., 2023). While a CNV was found in the cluster, it did not include *Cyp6aa1* and was not associated with resistance to deltamethrin. In Cameroon, a non-CNV haplotype has been shown to be associated with pyrethroid resistance (Kengne-Ouafo et al., 2024), while two large signals of selection are found around the same gene cluster, in the absence of any CNVs (*Anopheles gambiae* 1000 Genomes Consortium et al., 2017). While these haplotypes have not yet been phenotypically tested, we believe it very likely that it is associated with deltamethrin resistance, given the consistent results coming out of our study and the wider literature.

Resistance mutations in the deltamethrin target site, *Vgsc*, were very rare in our data, with only 2 samples in Moshi carrying the *Vgsc*-995F mutation. Strikingly, these two *Vgsc*-995F haplotypes were of different evolutionary origins, and have not introgressed into *An. arabiensis* from *An. gambiae. Vgsc*-995F has been consistently present but rare in Moshi over the 10 years preceding our collections (Kulkarni et al., 2006; Matowo et al., 2014). Our results suggest that the mutation has independently originated twice in *An. arabiensis*, and been under enough selective pressure to persist in the population, but not to reach high frequency, despite pyrethroid-driven evolution evidenced by the presence of P450-based metabolic resistance. One possibility is that the benefits of target site resistance to pyrethroids are lower in *An. arabiensis* than in *An. gambiae* s.s. and *An. coluzzii*, or that the physiological costs of such resistance are higher. However, the high frequency of *Vgsc*-995 mutations in *An. arabiensis* from West Africa suggests that target site resistance can be maintained in this species. The explanation for these differences may lie in the evolutionary history of these populations, their past exposure to DDT (which has the same target site, but different metabolic resistance pathways) and the order in which target site and metabolic resistance first appeared in the population.

Our agnostic genome-wide scans also revealed an association with deltamethrin resistance around three other detoxification loci: two cytochrome P450 clusters (*Cyp4h16-Cyp4h18* and *Cyp12f1*-*Cyp12f4*) and a glutathione-S-transferase cluster (*Gstd*). *Cyp4h17*, a member of the first P450 cluster, was highlighted as one of the most strongly upregulated genes in a genome-wide meta-analysis of resistant *Anopheles* expression data (Ingham and Nagi, 2024), suggesting at a role in resistance, which our data indicate could be against deltamethrin. In the *Cyp12f* cluster, *Cyp12f2* and *Cyp12f3* showed allelic imbalance in gene expression in F1 crosses between resistant and susceptible colonies of *An. gambiae*, suggesting differential *cis* regulation of expression linked to resistance (Dyer et al., 2023). Furthermore, the presence of *Cyp12f1* CNVs in both *An. gambiae* and *An. coluzzii* also hints at a possible role of this gene in resistance. When we originally described CNVs genome-wide in these two species (Lucas et al., 2019), we listed all cytochrome P450s, glutathione-S-transferases and carboxylesterases in which a CNV had been found. All of the genes in this list are in gene clusters that had previously, or have since, been shown to play a role in insecticide resistance in *Anopheles*, with the largest exception being *Cyp12f1*, which was only known for the possible association of *Cyp12f* genes with bendiocarb resistance in *An. gambiae* from Cameroon (Antonio-Nkondjio et al., 2016). It appears that all the metabolic genes in which we had identified CNVs have now accumulated evidence of association with resistance, suggesting that the presence of CNVs in such genes should in and of itself be considered as likely predictive evidence of a role in insecticide resistance.

As with *Cyp4h17, Gstd3* was also highlighted as consistently differentially expressed in resistant field populations compared to laboratory colonies in a transcriptomic meta-analysis (Ingham and Nagi, 2024), and we further showed equivocal evidence of an association of copy number of this gene with resistance to deltamethrin in our data. Further evidence is needed to determine the importance of this gene in resistance, and the insecticides to which resistance is most strongly conferred.

A conspicuous absence of signal in our data was in the region of *Cyp9k1* on chromosome X, which in *An. gambiae* and *An. coluzzii* showed evidence of association to both deltamethrin and PM (Main et al., 2015; Vontas et al., 2018; Hearn et al., 2022).

We have suggested that, given the challenge of genetically monitoring such a diverse landscape of resistance mutations in this cluster, more general methods, such as measuring gene expression directly, should be researched and developed to complement genetic screening panels (Lucas et al., 2023). We add to this the suggestion that including a measure of copy number in *Cyp6aa1* in resistance monitoring panels would present an alternative solution which, while being less encompassing, would perhaps come with fewer challenges.

Our results also provide evidence of resistance to PM in *An. arabiensis* from Tanzania. We found resistance in only one site, Moshi, while in Muleba there was full susceptibility, despite PM-based IRS having been applied there three years before our sampling (Protopopoff *et* al., 2018). Given the predominance of farming in Moshi, exposure to agricultural pesticides may be the cause of the elevated resistance to PM in this region. This exposure may have been more pervasive, over a longer period of time, and may therefore have been more effective at driving the evolution of resistance than IRS in outdoor-biting species such as *An. arabiensis*.

## 4. Methods

### 4.1 Sample collection and resistance characterisation

Mosquito larvae were collected from June to August 2018 from two locations in Tanzania, Moshi [-3.384, 37.349] and Muleba [-2.092, 31.574] (Supplementary Data S1). Moshi is an irrigated agricultural area south of Mount Kilimanjaro where *An. arabiensis* has historically been the primary vector of malaria (Matowo et al., 2014). Bednet distribution campaigns may have created selective pressure, although resistance levels to pyrethroids are generally moderate (Matowo et al., 2014). Muleba is on the border of lake Victoria and has been the site of IRS campaigns since 2007, and of vector control trials involving bednets and IRS. The village in which we collected our samples (Kyamyorwa) was targeted with lambda-cyhalothrin IRS from 2007 to 2011, bendiocarb IRS in 2011-2012 (Matowo et al., 2015) and PM IRS from 2014 to 2017 (Charlwood et al., 2018). Historically, *An. gambiae s*.*s*. has been the dominant malaria vector in the region, but the intervention trials have resulted in a large reduction in numbers for that species, and the preponderance of *An. arabiensis* (*Charlwood et al*., *2018)*.

Mosquito larvae collected from Moshi and Muleba were respectively transported for rearing to the insectary at the National Institute for Medical Research (NIMR), Amani Centre and the NIMR Mwanza insectary. 3-5 day-old females were characterised for resistance to insecticides (deltamethrin or PM) using our previously described method (Lucas et al., 2023). Briefly, we first performed a dose-response experiment to establish lethal doses to each insecticide in each location, and then identified susceptible mosquitoes as ones that were killed by a relatively low dose of insecticide, and resistant mosquitoes as ones surviving a relatively high dose (Fig. 3). This created greater phenotypic separation between susceptible and resistant samples, and thus greater power to detect significant associations. The results of the dose-response experiment, the doses used for each insecticide / location, sampling locations and dates, list of specimens and molecular species identification (Santolamazza et al., 2008; Chabi et al., 2019) are available in Supplementary Data S1.

**Figure 3:**
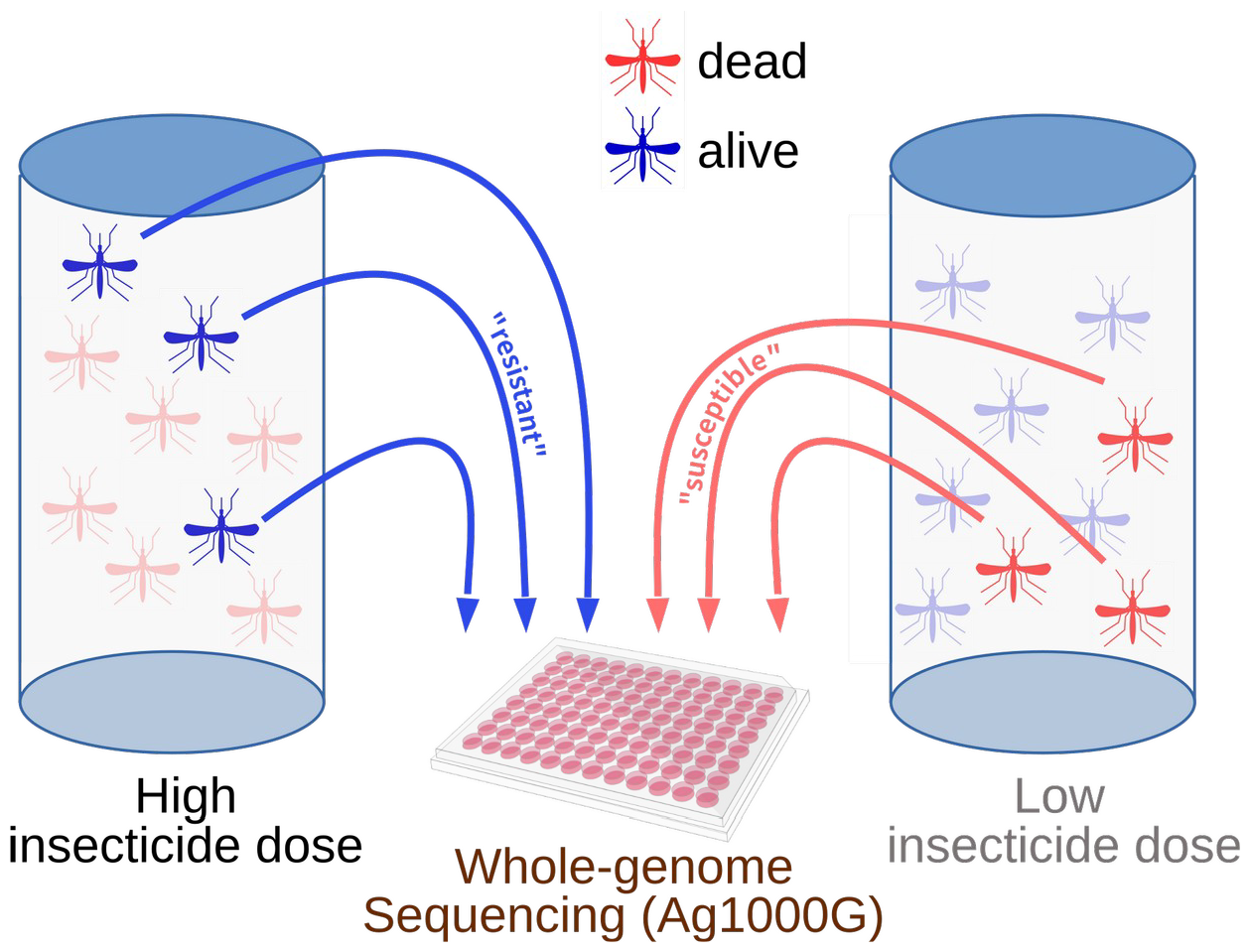
Summary of phenotyping protocol to obtain good separation of resistant and susceptible phenotypic groups for whole genome sequencing.

### 4.2 Whole genome sequencing and bioinformatic analysis

Overall, 489 samples were sequenced by the Ag1000G (full details of the pipeline: https://malariagen.github.io/vector-data/ag3/methods.html). Sample QC removed 2 samples for cross-contamination (*alpha* > 4.5%), 15 samples for low coverage (coverage < 10x, or less than 50% of genome with coverage > 1x), 4 samples as apparent technical replicates and 1 sample for unclear sex calling. 467 samples passed QC filtering. Data were released as part of Ag1000G release 3.7. All analyses using SNPs were performed using only loci that passed Ag1000G site quality filters.

We investigated CNVs in six regions with previously known association with insecticide resistance (*Ace1, Cyp6aa/Cyp6p, Cyp6m/Cyp6z, Gste, Coeae1f/Coeae2f*). We used diagnostic reads associated with known CNV alleles (https://www.malariagen.net/data, (Nagi et al., 2024) to identify these alleles in our data. To detect novel CNV alleles in these regions and in *Coeaexg*, we used the Ag1000G coverage-based CNV data, which applies a hidden Markov model (HMM) to normalised sequencing coverage to estimate the copy number state in 300 bp genomic windows (Lucas et al., 2019). In the *Cyp6aa/Cyp6p, Coeaexf* and *Coeaexg* regions, the HMM indicated the presence of CNVs in samples without known CNV alleles. We described these CNVs by manually identifying consistent discordant reads (reads mapping facing away from each-other in the reference genome) and soft-clipped reads matching the start and end points of coverage changes. These diagnostic reads will allow detection of these alleles in other whole genome sequencing datasets. The new CNVs have now been integrated into the Ag1000G CNV screening pipeline, and details of their start and end points can be found in Supplementary Data S2.

Copy number of individual genes was calculated as the mode of the HMM state within each gene. In the case of CNVs in *Coeaexf*, copy number often far exceeded the maximum copy number state allowed in the HMM (10 extra copies). To obtain a more accurate value of copy number for this CNV, we instead took the median raw normalised coverage for all windows found within the CNV region (positions 37282000 to 37295000) and subtracted 2 (the normal diploid copy number).

The KING statistic of kinship (Manichaikul et al., 2010) was calculated using NGSRelate (Hanghøj et al., 2019) using genome-wide SNPs, excluding regions of genomic inversions (2L:13-38Mb and 2R:19-33Mb). We used a threshold KING value of 0.185 (Supplementary Methods) to classify full sibs. From each sib group, we randomly chose a single individual to retain for all analyses described below, discarding the others. The exception to this was the calculation of F_ST_, where it was computationally feasible to permute which sibs were removed (see below).

Selection scans were performed using H_12_ (Garud et al., 2015) and H_1X_ (Miles, 2021). H_1X_ is a measure of haplotype sharing between two populations, calculated as 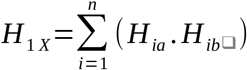, where *n* is the number of haplotypes found in either population, and *H*_*ia*_ and *H*_*ib*_ are the frequencies of haplotype *i* in populations *a* and *b* respectively. High values of H_1X_ indicate that high frequency (ie: swept) haplotypes are shared between the two populations.

### 4.5 Phenotypic association of CNVs and known resistance SNPs

We investigated association between resistance phenotype and individual genetic markers (CNVs or SNPs) using generalised linear models implemented in R v4 (R Core Team, 2021), with binomial error and a logit link function, with phenotype as the dependent variable and genotypes as independent variables. SNP genotypes were coded numerically as the number of mutant alleles (possible values of 0, 1 and 2), CNV alleles were coded as presence / absence, and gene copy number was coded as the number of extra copies. Starting from the null model, we proceeded by stepwise model building, adding the most highly significant marker at each step until no remaining markers provided a significant improvement.

To calculate the statistical power of finding an effect of *Coeae1f* on resistance to PM, we took the data in which a significant association had previously been found (Nagi et al., 2024) and calculated that mortality had been 44.2% in wild-type individuals and 16.7% in individuals carrying a CNV. Mortality in wild-type individuals was almost the same in Moshi (44.4%). We therefore ran 1000 Monte Carlo simulations using our sample size and CNV frequency from Moshi, with the mortalities observed in Ghana. For each randomisation, we ran the same glm as on the real data, and calculated the proportion of simulations in which we observed *P* < 0.05.

### 4.6 Windowed measures of differentiation (F_ST_, PBS, H_12_)

We calculated F_ST_ using the *moving_patterson_fst* function in *scikit-allel* (Miles and Harding, 2016) in a moving window of 1000 SNPs, after filtering SNPs for missing data and removing singletons. In order to take advantage of the full sample set despite non-independence of full siblings, we performed permutations in which one randomly chosen individual per sib group was used in the calculation of F_ST_. In Muleba, we performed 100 such permutations and calculated the mean F_ST_ of all permutations. In Moshi, the PM sample set contained no sibs, while the deltamethrin sample set contained only one pair of sibs and thus needed only averaging the two calculations of F_ST_ (removing each sib in turn).

Provisional windows of interest (“peaks”) were identified as ones with positive F_ST_ values three times further from the mode than the smallest negative value (Lucas et al., 2023). We then removed peaks that might be the result of the presence of a selective sweep, as opposed to true association of that sweep with resistance (Lucas et al., 2023), using Monte Carlo simulations creating 500 permutations of the phenotype labels and recalculating F_ST_ for each permutation. We retained peaks whose observed F_ST_ was greater than 99% of the simulations.

F_ST_ indicates any genetic differences between resistant and susceptible samples, but we expect that differences associated with resistance would be associated with the presence of swept haplotypes at higher frequency in the resistant compared to susceptible samples. To further filter the F_ST_ peaks, we therefore explored the presence of swept haplotypes within each peak. Haplotype clusters were determined by hierarchical clustering on pairwise genetic distance (Dxy) between haplotypes, and cutting the tree at a height of 0.001. Clusters comprised of at least 20 haplotypes were tested for association with phenotype using a generalised linear model with binomial error and logit link function, with phenotype as the response and sample genotype (number of copies of the haplotype) as a numerical independent variable. Peaks were discarded if they did not contain a haplotype positively associated with resistance.

We calculated H_12_ using the *garuds_h* function in *scikit-allel* in a moving window of 1000 SNPs, using phased biallelic SNPs. The ΔH_12_ metric was obtained by subtracting H_12_ in the susceptible samples from H_12_ in the resistant samples, with a positive value thus indicating a higher frequency of swept haplotypes in resistant samples. PBS between susceptible and resistant samples was calculated using segregating SNPs in 1000 SNP windows using the *pbs* function in *scikit-allel*, with *An. arabiensis* samples from Malawi, collected in 2015, as the outgroup (Ag1000G phase 3.0 data release). As with ΔH_12_, positive signals of PBS indicate stronger positive selection in the resistant samples. We identified provisional peaks in PBS by taking windows with a PBS value higher than 3 times the 95th centile of the PBS distribution. Using this threshold for H_12_ resulted in a very large number of provisional peaks across the entire genome, and we thus used 3 times the 98th centile as a threshold instead for H_12_. For both H_12_ and PBS, 500 monte carlo permutations of phenotype were performed as above to remove false positive peaks caused by the presence of swept haplotypes.

### 4.8 Genome-wide association analysis (GWAS)

We performed SNP-wise GWAS using SNPs with no missing data and a minor allele count of at least 5. In a previous study, we found that contamination of our samples by *Asaia* bacteria caused artefacts in our association analysis (Lucas et al., 2023). We therefore used Bracken (Lu et al., 2017) to estimate the amount of *Asaia* contamination in each sample and excluded SNP loci where genotype was correlated with *Asaia* levels (*P* < 0.05).

For each SNP, we used a GLM with binomial error and logit link function to obtain a *P* value of association for phenotype against genotype (coded as the number of non-reference alleles). We used *fdrtool* (Klaus and Strimmer, 2015) to perform false-discovery rate correction, with *Q* value threshold of 1%. We also used these data to perform a windowed analysis, identifying the 1000 most significant SNPs in each sample set and looking for 100,000bp windows that contained at least 10 SNPs among the top 1000.

## Availability of data and materials

Code used to analyse the data can be found in the github repository https://github.com/vigg-lstm/GAARD_east. Sequencing, alignment, SNP and CNV calling was carried out as part of the *Anopheles gambiae* 1000 genomes project v3.7 (https://www.malariagen.net/data). Raw reads were submitted to the ENA archive, and accession numbers are provided in Supplementary Data S1. Where we included data from West African samples, these formed part of ag1000G release v3.2, and the specific data that we used were drawn from the github repository https://raw.githubusercontent.com/vigg-lstm/GAARD_work/v2.0.

## Acknowledgements

This work was supported by the National Institute of Allergy and Infectious Diseases (NIAID R01-AI116811 to M.J.D. and D.W.) and the Medical Research Council (MR/T001070/1 to M.J.D., D.W. and E.R.L., MR/ P02520X/1 to M.J.D. and D.W.). The latter grant is a UK-funded award and is part of the EDCTP2 programme supported by the European Union. M.J.D. is supported by a Royal Society Wolfson Fellowship (RSWF\FT \180003). N. D. is supported by a Royal Society Dorothy Hodgkin Fellowship (DHF\R1\231087). We thank Charles Kayamba and Mathias Stephano for their assistance in mosquito sample collection, rearing and susceptibility testing. We also thank the *Anopheles gambiae* 1000 genomes project for carrying out the sequencing, quality control, SNP calling and for haplotype phasing the sequencing data, and Luciene Salas Jennings and Andrew Carey for providing administrative support to the project.

## Author contributions

B.B., J.E., S.D. and J.C. performed sample collection, rearing and bioassay testing. B.K., W.K. and A.E.-Y. oversaw sample collection, rearing and bioassay testing. E.J.R., D.P. and A.E.V.H. performed the lab work. A.M. oversaw the sample sequencing. A.M., N.J.H., and C.S.C performed sequence alignment, SNP calling and haplotype phasing. E.R.L. performed the CNV calling, genome scans, association studies and haplotype clustering. S.C.N. performed the kinship analyses. D.W., M.J.D., B. K. and A.E.-Y. designed the study. E.R.L., S.C.N., N.A.D., M.J.D. and D.W. wrote the manuscript. All authors read and approved the final manuscript.

