## Supplementary Figures and Tables for "Copy number variants underlie the major selective sweeps in insecticide resistance genes in *Anopheles arabiensis* from Tanzania"

### Electronic Supplementary Material
### Supplementary figures and tables

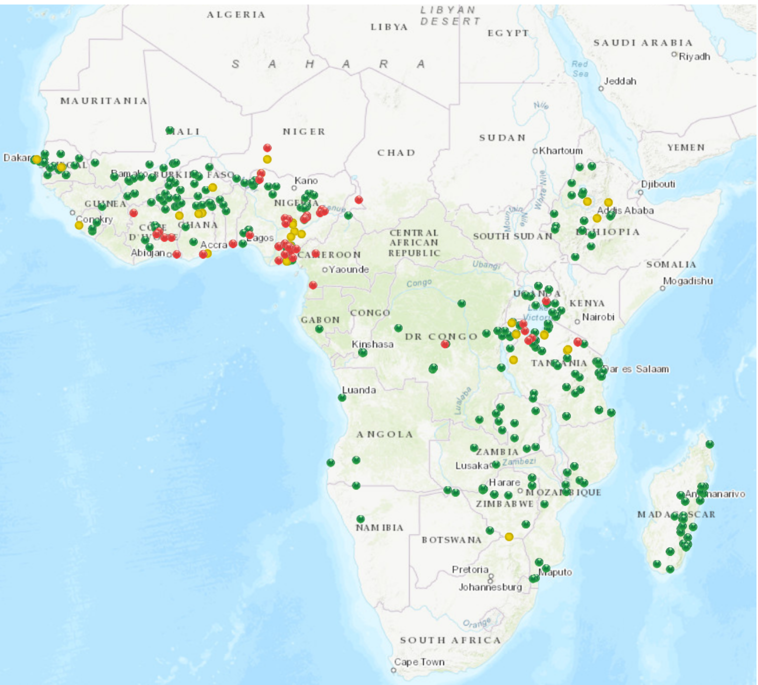


**Fig. S1**: Distribution of resistance to pirimiphos methyl detected by studies in *An. gambiae* sl over the last 10 years (2014-2024), indicating confirmed resistance (red), possible resistance (yellow) and susceptibility (green). Data obtained from IR-Mapper v2.0 `https://anopheles.irmapper.com/` on 29th of January 2024.

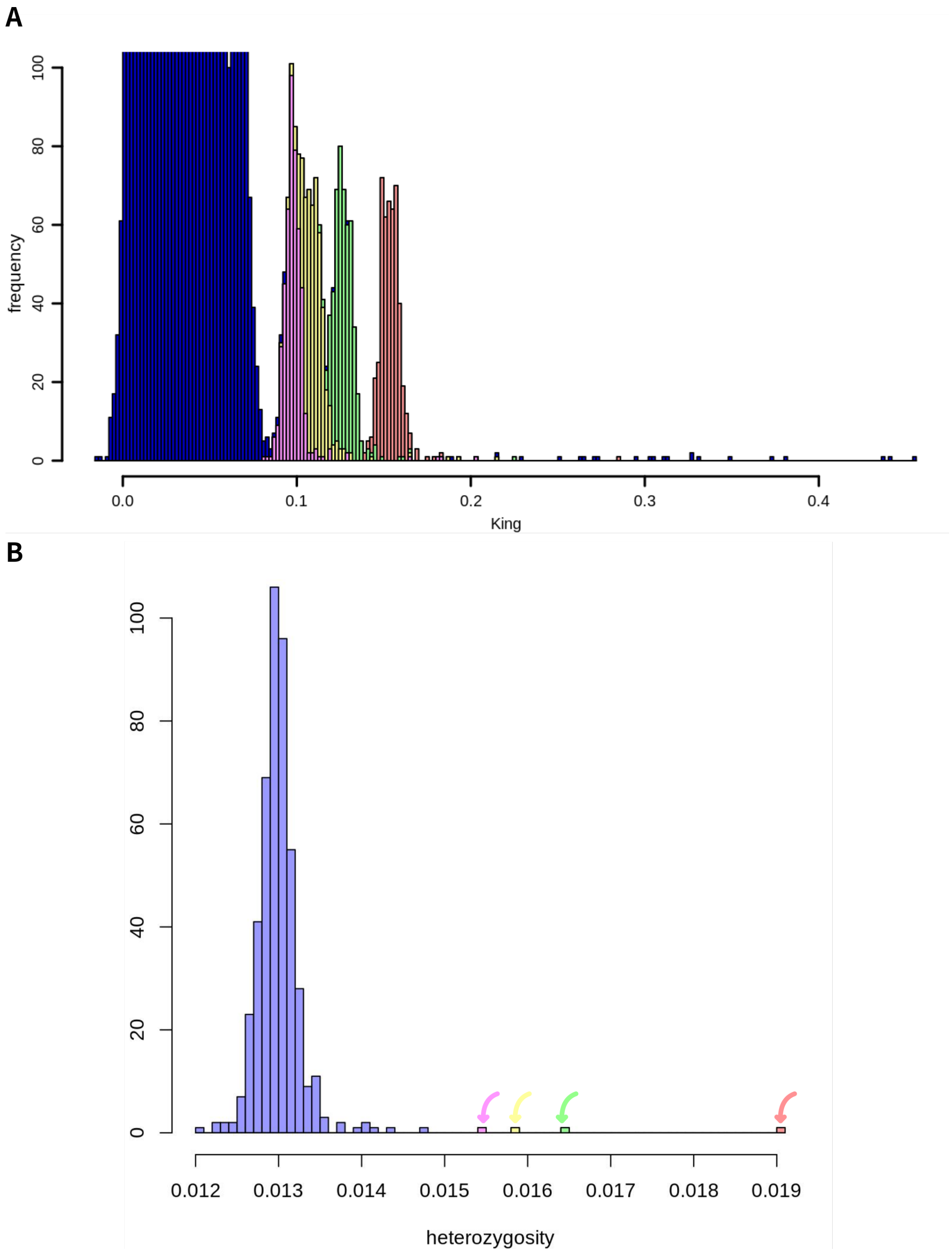


**Fig. S2**: **A** Histogram (zoomed in on the x axis to show right-hand tail) of all pair-wise KING relatedness values across all samples in our dataset. Four samples had universally high relatedness values to all samples. The relatedness values attributables to these four samples have been respectively coloured in pink, yellow, green and red on the histogram. **B** Histogram of heterozygosity values across all samples, showing that the same four samples (coloured and indicated with arrows) have elevated heterozygosity. As a result, these four samples had aritificially augmented KING values.

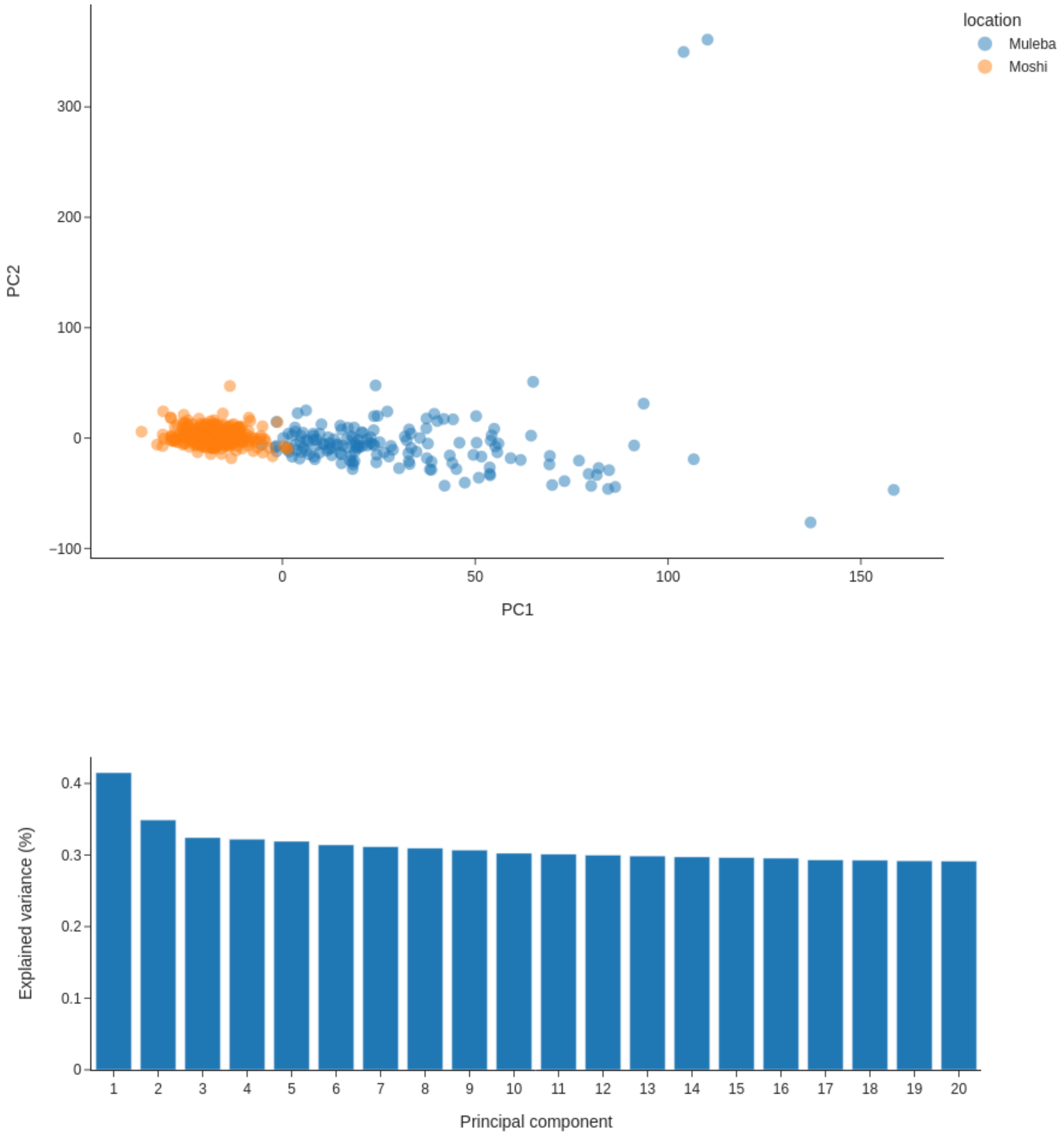


**Fig. S3**: PCA (using quality-filtered biallelic SNPs from genomic region 3L:15,000,000-41,000,000, euchromatic and free of chromosomal inverstions). Top panel shows clustering of samples by region. Bottom panel show variance explained by the first 10 PCs, indicating that PCs 3 onwards explain similar levels of variance and are thus likely only capturing noise.

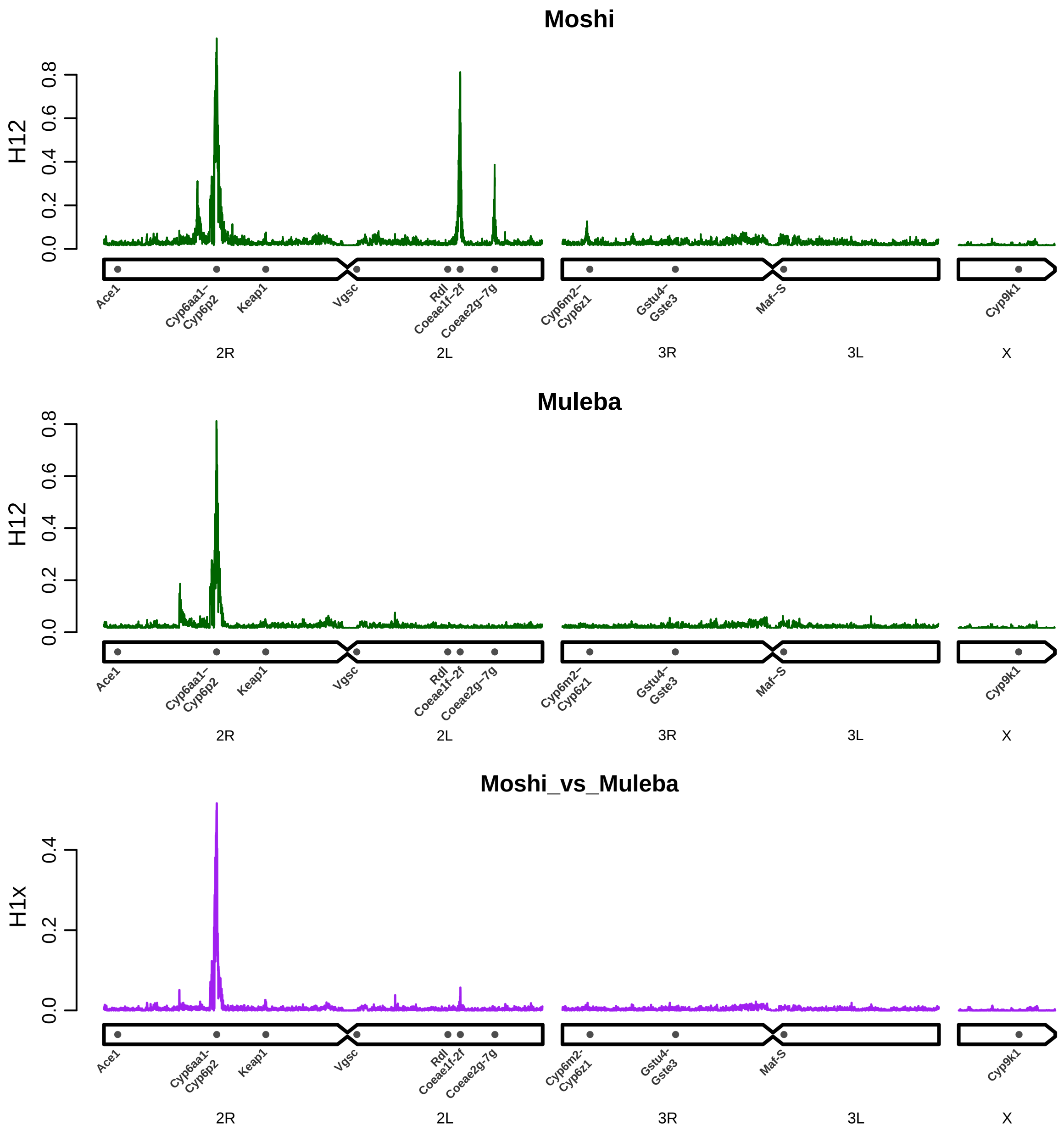


**Fig. S4**: Selection scans showing genome-wide H12 signal in Moshi (top) and Muleba (middle), as well as shared signals of selection (H1x) between the two sites (bottom).

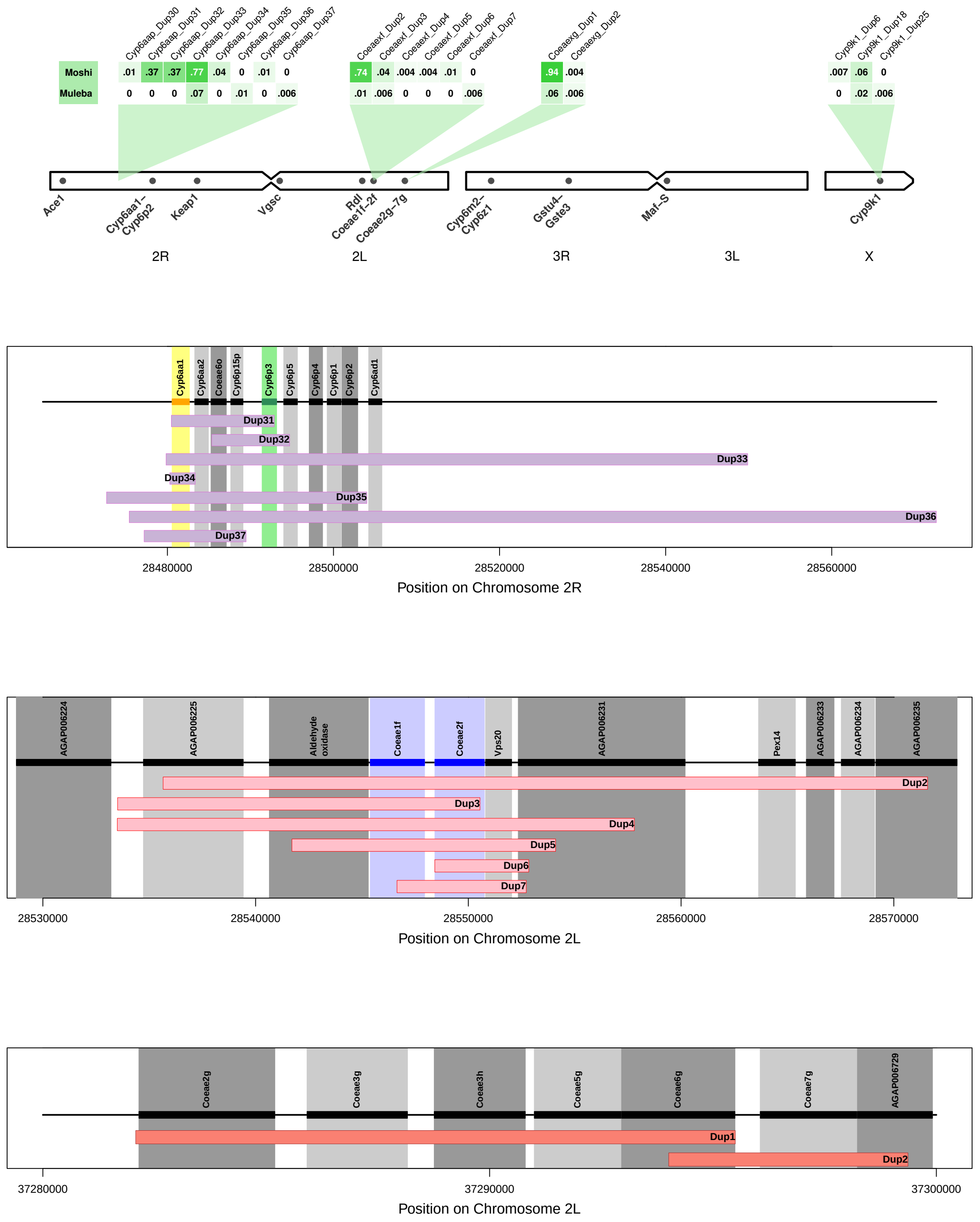


**Fig. S5**: **Top panel**: Frequency (proportion of samples carrying at least one copy) of known CNV alleles detected using diagnostic reads around *Ace1*, the *Cyp6aa / Cyp6p* cluster, *Gstd3*, the *Coeaexf* cluster, the *Coeaexg* cluster, the *Cyp6m / Cyp6z* cluster, the *Gste* cluster and *Cyp9k1*. Only CNV alleles with frequency > 0% are shown. Cell darkness provided as a visual aid for the magnitude of the value in each cell. **Bottom three panels**: genomic ranges of newly characterised CNV alleles, with genes previously shown to be important in resistance highlighed in colour. The precise genomic coordinates of each CNV allele can be found in Supplementary Data S2. The CNV in *Coezexg* in West African *An. coluzzii* is not shown as its start and end points are unknown.

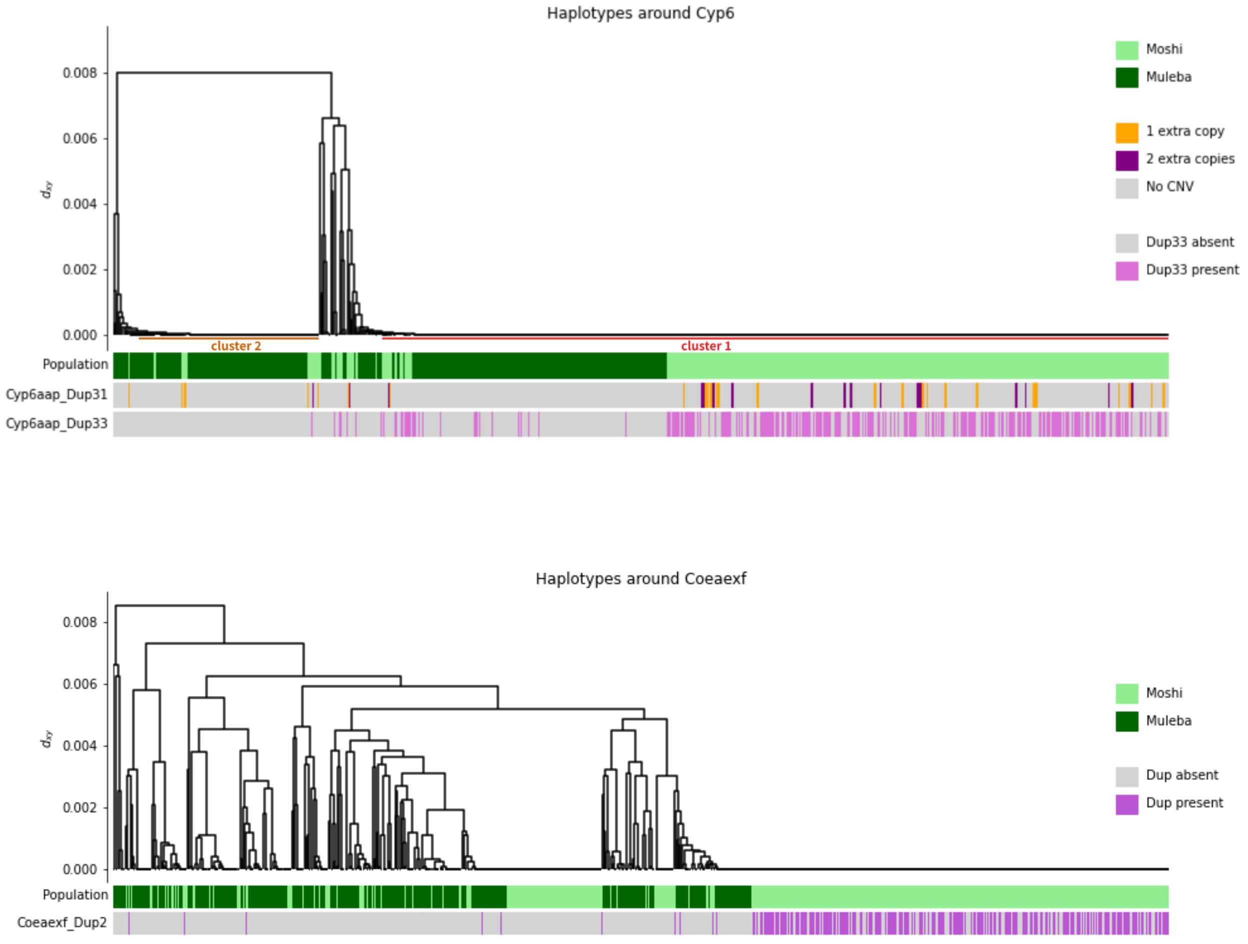


**Fig. S6**: Haplotype clustering of the genomic region around *Cyp6aa/Cyp6p* (top panel) showing that nearly all haplotypes belong to one of two selective sweeps. The less common cluster (cluster2) is predominantly found in Muleba and is not associated with CNVs. The more common cluster (cluster1), is found in both regions. Both *Cyp6aap*_Dup31 and *Cyp6aap*_Dup33 form a subset of haplotypes in cluster1. For *Cyp6aap*_Dup33, it was possible to assign presence (mauve) or absence (grey) of the CNV for each haplotype. For *Cyp6aap*_Dup31, it was only possible to determine whether the mosquito to which the haplotype belongs had a single extra copy (yellow), two (purple) or none (grey). A single extra copy indicates the sample is heterozygous for the CNV, and thus haplotypes labelled in yellow may not themselves carry the CNV. Similarly, in the *Coeaexf* region (bottom panel), haplotypes bearing the CNV allele *Coeaexf*_Dup2 represented a subset of haplotypes from a large swept cluster. Clustering was performed using 500 SNPs in each region, and CNV alleles were phased by identifying SNPs that were highly correlated with their presence / absence.. Full workings to reproduce this analysis can be found at `https://github.com/vigg-lstm/GAARD_east/blob/main/CNV_analysis/sweeps`.

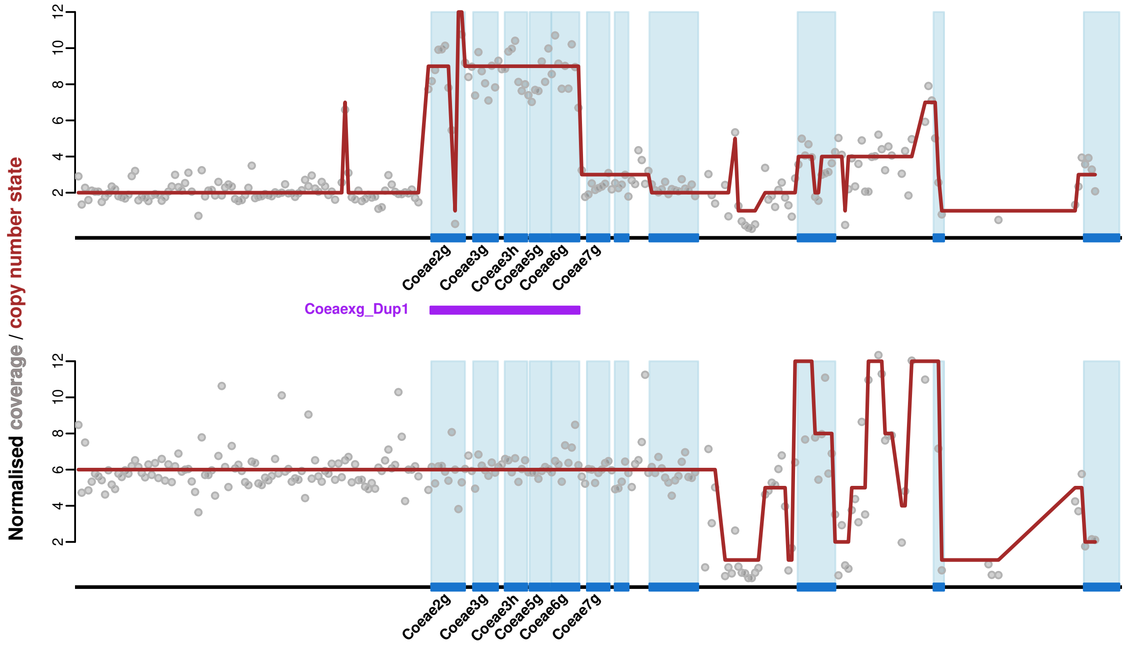


**Fig. S7**: Example coverage traces for CNVs in *Coeaexg*. Grey points show raw coverage (in 300 bp windows) normalised to a copy number of two (normal diploid copy number); brown line shows the output of the Hidden Markov Model (HMM) through the coverage data, indicating the predicted copy number state in each window. The genomic region to the right of the *Coeaexg* cluster has erratic coverage, suggesting a repeat region or poor genome assembly. CNVs in *Coeaexg* are evident from the HMM being consistently above the normal value of 2 across the region. The top plot shows an example sample carrying the main CNV allele found in our dataset of *An. arabiensis* from Tanzania (*Coeaexg*_Dup1, region coveraged by the CNV shown by purple bar). The bottom plot shows an example sample of *An. coluzzii* from Korle-Bu (Ghana), where the CNV extends far to the left, and into the region of erratic coverage to the right, thus meaning that we could not identify discordant reads that could tag individual CNV alleles in this population.

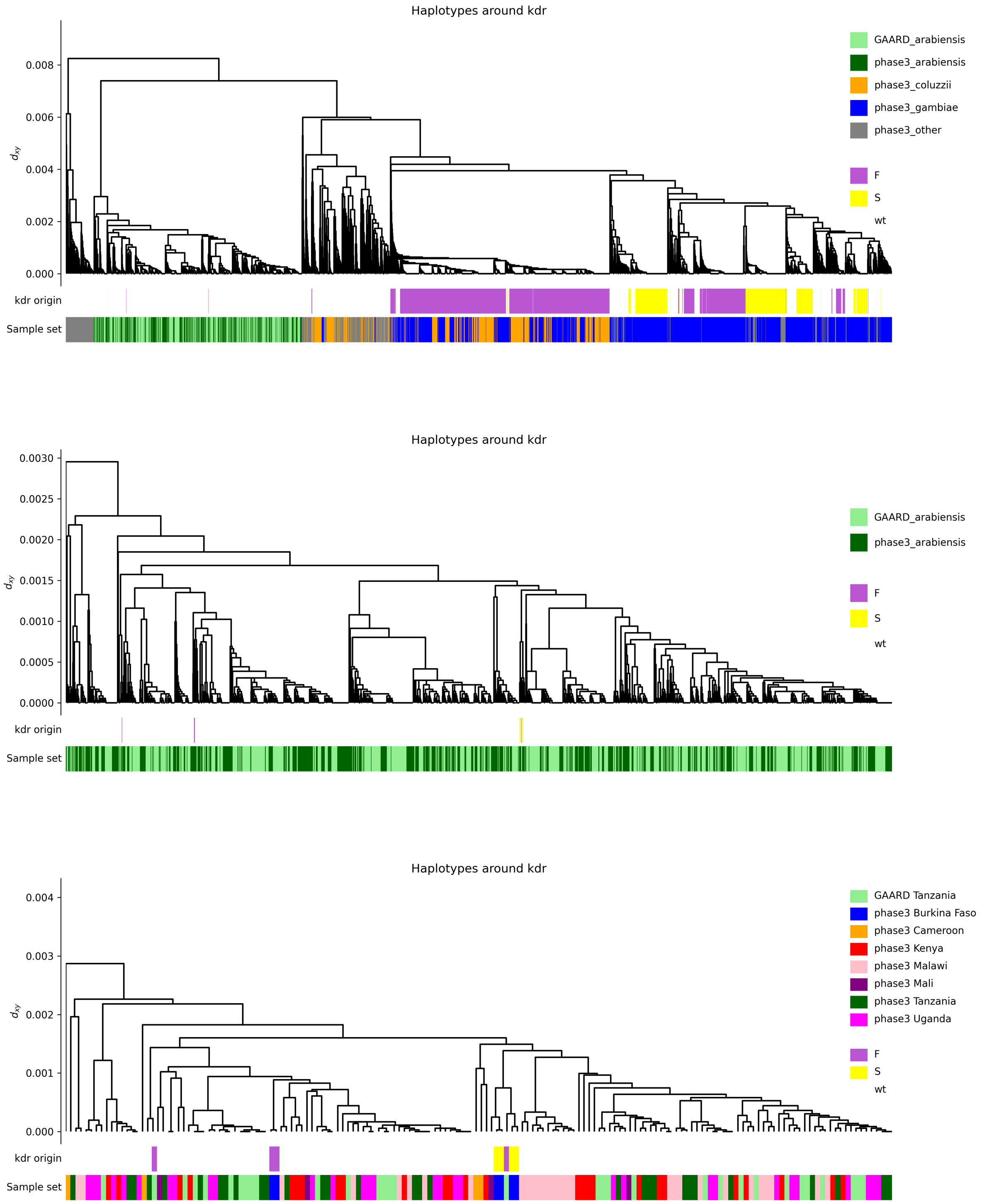


**Fig. S8**: Clustering of haplotypes around the *Vgsc* genomic region (2L:2358158-2431617) reveals a diversity of *Vgsc*-995 origins in *An. arabiensis*, none of which are introgressed from *An. gambiae* or *An. coluzzii*. Combining our data with all haplotypes from phase 3 of Ag1000G (top) shows *An. arabiensis* haplotypes forming their own cluster, distinct from other species. When keeping only *An. arabiensis* haplotypes (middle), three different *Vgsc*-995F clusters are seen, despite only four such haplotypes exising in the dataset. The bottom plot shows all eight *Vgsc*-995 mutant haplotypes (two from our Tanzanian data, six from phase3 samples from Burkina Faso) and a random sub-sample of wild-type *An. arabiensis*, allowing a closer view of sample set labels for interpretation. The two Tanzanian *Vgsc*-995F haplotypes appear to be independent origins, one of which clusters more closely with haplotypes from Burkina Faso than with other Tanzanian haplotypes.

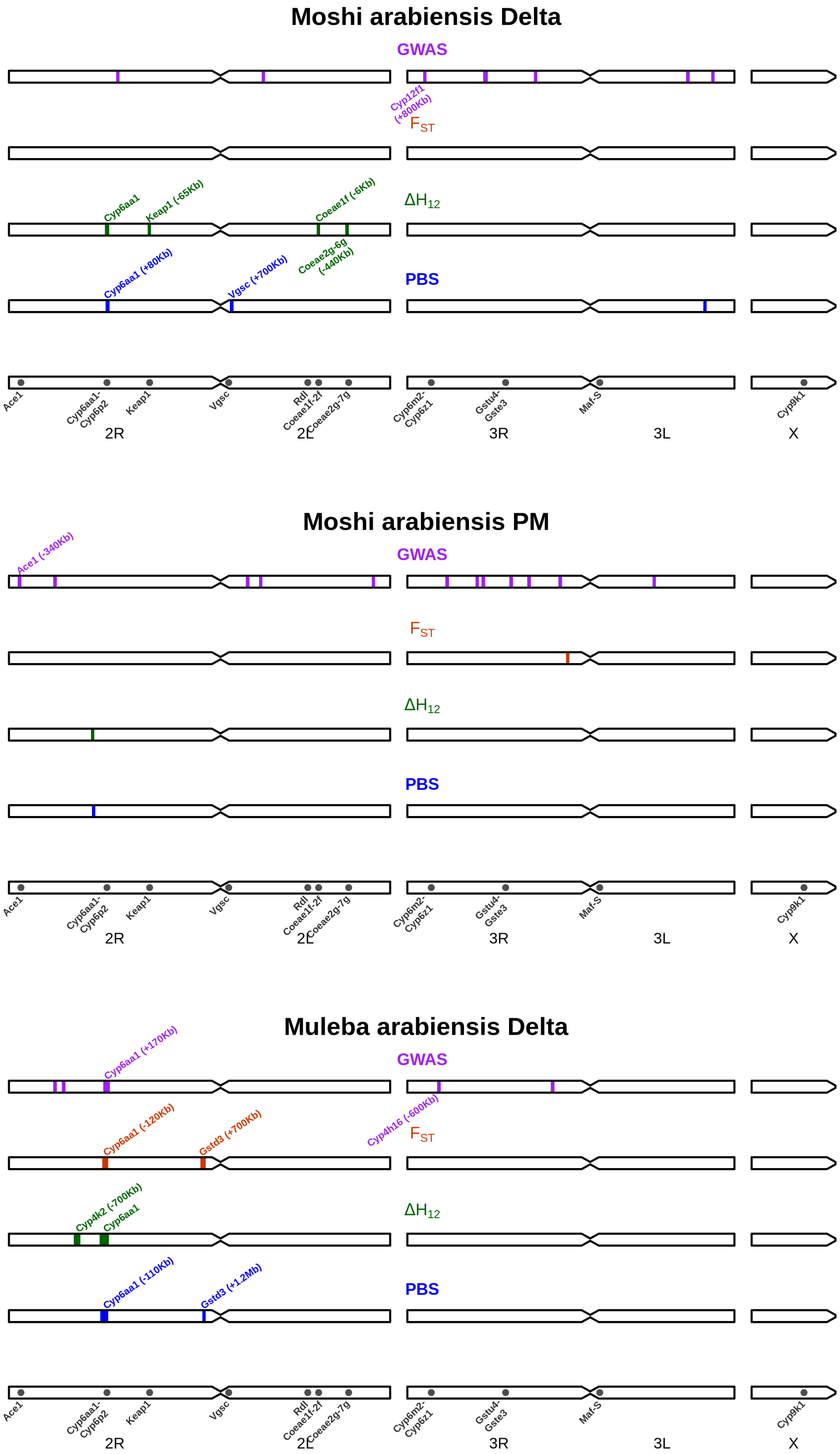


**Fig. S9**: Genomic regions implicated in insecticide resistance by each of our four approaches. For the global GWAS method, these are 100,000 bp windows which contained at least 10 of the top 1000 significant SNPs. For $F_{ST}$, these are significant peaks which contained at least one haplotype significantly positively associated with resistance (Supplementary Data S2). For $\Delta H_{12}$ and PBS, these are significant positive peaks (ie: indicating stronger signals of selection in resistant compared to susceptible samples). Regions are annotated with genes discussed in the manuscript as possibly causing the signal. Genomic distances in brackets indicate the distance of the peak either to the left (-) or right (+) of the gene in question.

**Table S1**: Non-synonymous SNPs in the *Ace1* gene with a minor allele count (MAC) of at least 5 (out of 302 haplotypes) in the Moshi PM dataset. *P* value is the result of a logistic regression of genotype vs phenotype for each SNP. Effect direction indicates whether samples carrying the SNP tended to be more ("Resistant") or less ("Susceptible") likely to survive insecticide exposure. None of the five SNPs were associated with resistance.

| **Chrom** | **Pos** | **MAC** | ***P*** | **Effect direction** | **Nucleotide change** | **Amino acid change** |
|---|---|---|---|---|---|---|
| 2R | 3489390 | 6 | 0.83 | Susceptible | 178G>A | Ala60Thr |
| 2R | 3489397 | 11 | 0.54 | Susceptible | 185T>G | Val62Gly |
| 2R | 3493415 | 6 | 0.29 | Susceptible | 1948G>A | Glu650Lys |
| 2R | 3493739 | 7 | 0.88 | Resistant | 2165T>C | Ile722Thr |
| 2R | 3493765 | 5 | 0.79 | Resistant | 2191G>A | Ala731Thr |
