## Supplementary Data S6 for "Copy number variants underlie the major selective sweeps in insecticide resistance genes in *Anopheles arabiensis* from Tanzania": Supplementary_Data_S6.html

gwas\_summary


### GAARD GWAS results summary

Each plot shows the 1000 most significant SNPs in the GWAS for that
sample set. At the top, the P value for each SNP is shown, colour-coded
by chromosome. Underneath is a heatmap showing the pairwise correlation
between all 1000 SNPs, helping to see groups of SNPs that are in linkage
disequilibrium. Each point in the *P*-value plot is lined up with
its column in the heatmap. Beneath the column, lines heatmap columns to
a map of the genome to show the genomic position of each SNP. Black
horizontal bars between the heatmap and the linking lines indicate
genomic regions containing a high density of these top 1000 SNPs (100 kb
windows that contained at least 10 SNPs).

---

Moshi\_*arabiensis*\_Delta  
Muleba\_*arabiensis*\_Delta  
Moshi\_*arabiensis*\_PM

---

#### Moshi\_*arabiensis*\_Delta

---

#### Muleba\_*arabiensis*\_Delta

---

#### Moshi\_*arabiensis*\_PM

---
