## Supplementary Data S5 for "Copy number variants underlie the major selective sweeps in insecticide resistance genes in *Anopheles arabiensis* from Tanzania": Supplementary_Data_S5.html

window\_PBS\_summary


### PBS windows of interest

For each sample set, we first provide a summary plot of PBS across
the genome, with PBS shown in blue and the results of the 200
randomisations shown behind in grey. Windows identified as peaks are
highlighted by points, colour-coded by whether they are significantly
higher than expected based on the simulations (green) or not (purple).
For each significant window (green points), we then provide its own plot
showing the significant (*P* < 0.01) SNPs found in the region
of that window, and their -log10(Pvalue) of association with phenotype.
Red points indicate non-synonymous SNPs, blue points indicate all other
SNPs. Point shape indicates whether the mutant allele at that SNP is
associated with increased (circle) or decreased (triangle) resistance.
Dark points in the centre of the plot show SNPs within the significant
window, light points on the sides show SNPs in the region 10,000 bp
either side of the window.

Legend  
Moshi\_*arabiensis*\_Delta  
Muleba\_*arabiensis*\_Delta  
Moshi\_*arabiensis*\_PM

---

#### Plot legend

---

#### Moshi\_*arabiensis*\_Delta

Moshi\_arabiensis\_Delta\_2L:3276066  
Moshi\_arabiensis\_Delta\_2R:28595065  
Moshi\_arabiensis\_Delta\_2R:28709379  
Moshi\_arabiensis\_Delta\_2R:28768446  
Moshi\_arabiensis\_Delta\_3L:33374272

##### Moshi\_arabiensis\_Delta\_2L\_3276066

##### Moshi\_arabiensis\_Delta\_2R\_28595065

##### Moshi\_arabiensis\_Delta\_2R\_28709379

##### Moshi\_arabiensis\_Delta\_2R\_28768446

##### Moshi\_arabiensis\_Delta\_3L\_33374272

---

#### Muleba\_*arabiensis*\_Delta

Muleba\_arabiensis\_Delta\_2R:27108686  
Muleba\_arabiensis\_Delta\_2R:27191090  
Muleba\_arabiensis\_Delta\_2R:27670025  
Muleba\_arabiensis\_Delta\_2R:27698574  
Muleba\_arabiensis\_Delta\_2R:27896706  
Muleba\_arabiensis\_Delta\_2R:27935028  
Muleba\_arabiensis\_Delta\_2R:28112399  
Muleba\_arabiensis\_Delta\_2R:28174246  
Muleba\_arabiensis\_Delta\_2R:28264928  
Muleba\_arabiensis\_Delta\_2R:28307853  
Muleba\_arabiensis\_Delta\_2R:28347791  
Muleba\_arabiensis\_Delta\_2R:56761889

##### Muleba\_arabiensis\_Delta\_2R\_27108686

##### Muleba\_arabiensis\_Delta\_2R\_27191090

##### Muleba\_arabiensis\_Delta\_2R\_27670025

##### Muleba\_arabiensis\_Delta\_2R\_27698574

##### Muleba\_arabiensis\_Delta\_2R\_27896706

##### Muleba\_arabiensis\_Delta\_2R\_27935028

##### Muleba\_arabiensis\_Delta\_2R\_28112399

##### Muleba\_arabiensis\_Delta\_2R\_28174246

##### Muleba\_arabiensis\_Delta\_2R\_28264928

##### Muleba\_arabiensis\_Delta\_2R\_28307853

##### Muleba\_arabiensis\_Delta\_2R\_28347791

##### Muleba\_arabiensis\_Delta\_2R\_56761889

---

#### Moshi\_*arabiensis*\_PM

Moshi\_arabiensis\_PM\_2R:24644689  
Moshi\_arabiensis\_PM\_2R:24668375

##### Moshi\_arabiensis\_PM\_2R\_24644689

##### Moshi\_arabiensis\_PM\_2R\_24668375

---
