## Supplementary Data S4 for "Copy number variants underlie the major selective sweeps in insecticide resistance genes in *Anopheles arabiensis* from Tanzania": Supplementary_Data_S4.html

window\_H12\_summary


### H12 windows of interest

For each sample set, we first provide a summary plot of *Δ*H_12_ (H_12_ in
susceptible samples subtracted from that in resistant samples) across
the genome, with *Δ*H_12_ shown in green and
the results of the 200 randomisations shown behind in grey. Windows
identified as peaks are highlighted by points, colour-coded by whether
they are significantly higher than expected based on the simulations
(green) or not (purple). For each significant window (green points), we
then provide its own plot showing the significant (*P* < 0.01)
SNPs found in the region of that window, and their -log10(Pvalue) of
association with phenotype. Red points indicate non-synonymous SNPs,
blue points indicate all other SNPs. Point shape indicates whether the
mutant allele at that SNP is associated with increased (circle) or
decreased (triangle) resistance. Dark points in the centre of the plot
show SNPs within the significant window, light points on the sides show
SNPs in the region 10,000 bp either side of the window.

Legend  
Moshi\_*arabiensis*\_Delta  
Muleba\_*arabiensis*\_Delta  
Moshi\_*arabiensis*\_PM

---

#### Plot legend

---

#### Moshi\_*arabiensis*\_Delta

Moshi\_arabiensis\_Delta\_2L:28513425  
Moshi\_arabiensis\_Delta\_2L:36831343  
Moshi\_arabiensis\_Delta\_2R:28385301  
Moshi\_arabiensis\_Delta\_2R:28568181  
Moshi\_arabiensis\_Delta\_2R:28621705  
Moshi\_arabiensis\_Delta\_2R:28662339  
Moshi\_arabiensis\_Delta\_2R:28708855  
Moshi\_arabiensis\_Delta\_2R:40849418

##### Moshi\_arabiensis\_Delta\_2L\_28513425

##### Moshi\_arabiensis\_Delta\_2L\_36831343

##### Moshi\_arabiensis\_Delta\_2R\_28385301

##### Moshi\_arabiensis\_Delta\_2R\_28568181

##### Moshi\_arabiensis\_Delta\_2R\_28621705

##### Moshi\_arabiensis\_Delta\_2R\_28662339

##### Moshi\_arabiensis\_Delta\_2R\_28708855

##### Moshi\_arabiensis\_Delta\_2R\_40849418

---

#### Muleba\_*arabiensis*\_Delta

Muleba\_arabiensis\_Delta\_2R:19368988  
Muleba\_arabiensis\_Delta\_2R:19709283  
Muleba\_arabiensis\_Delta\_2R:19865728  
Muleba\_arabiensis\_Delta\_2R:20325777  
Muleba\_arabiensis\_Delta\_2R:26914367  
Muleba\_arabiensis\_Delta\_2R:26935183  
Muleba\_arabiensis\_Delta\_2R:27029713  
Muleba\_arabiensis\_Delta\_2R:27071001  
Muleba\_arabiensis\_Delta\_2R:27087241  
Muleba\_arabiensis\_Delta\_2R:27101453  
Muleba\_arabiensis\_Delta\_2R:27116593  
Muleba\_arabiensis\_Delta\_2R:27134263  
Muleba\_arabiensis\_Delta\_2R:27187264  
Muleba\_arabiensis\_Delta\_2R:27202906  
Muleba\_arabiensis\_Delta\_2R:27218667  
Muleba\_arabiensis\_Delta\_2R:27252717  
Muleba\_arabiensis\_Delta\_2R:27385714  
Muleba\_arabiensis\_Delta\_2R:27867820  
Muleba\_arabiensis\_Delta\_2R:27971341  
Muleba\_arabiensis\_Delta\_2R:27994335  
Muleba\_arabiensis\_Delta\_2R:28037251  
Muleba\_arabiensis\_Delta\_2R:28061969  
Muleba\_arabiensis\_Delta\_2R:28128599  
Muleba\_arabiensis\_Delta\_2R:28200653  
Muleba\_arabiensis\_Delta\_2R:28232572  
Muleba\_arabiensis\_Delta\_2R:28335316  
Muleba\_arabiensis\_Delta\_2R:28366295  
Muleba\_arabiensis\_Delta\_2R:28439507  
Muleba\_arabiensis\_Delta\_2R:28616955

##### Muleba\_arabiensis\_Delta\_2R\_19368988

##### Muleba\_arabiensis\_Delta\_2R\_19709283

##### Muleba\_arabiensis\_Delta\_2R\_19865728

##### Muleba\_arabiensis\_Delta\_2R\_20325777

##### Muleba\_arabiensis\_Delta\_2R\_26914367

##### Muleba\_arabiensis\_Delta\_2R\_26935183

##### Muleba\_arabiensis\_Delta\_2R\_27029713

##### Muleba\_arabiensis\_Delta\_2R\_27071001

##### Muleba\_arabiensis\_Delta\_2R\_27087241

##### Muleba\_arabiensis\_Delta\_2R\_27101453

##### Muleba\_arabiensis\_Delta\_2R\_27116593

##### Muleba\_arabiensis\_Delta\_2R\_27134263

##### Muleba\_arabiensis\_Delta\_2R\_27187264

##### Muleba\_arabiensis\_Delta\_2R\_27202906

##### Muleba\_arabiensis\_Delta\_2R\_27218667

##### Muleba\_arabiensis\_Delta\_2R\_27252717

##### Muleba\_arabiensis\_Delta\_2R\_27385714

##### Muleba\_arabiensis\_Delta\_2R\_27867820

##### Muleba\_arabiensis\_Delta\_2R\_27971341

##### Muleba\_arabiensis\_Delta\_2R\_27994335

##### Muleba\_arabiensis\_Delta\_2R\_28037251

##### Muleba\_arabiensis\_Delta\_2R\_28061969

##### Muleba\_arabiensis\_Delta\_2R\_28128599

##### Muleba\_arabiensis\_Delta\_2R\_28200653

##### Muleba\_arabiensis\_Delta\_2R\_28232572

##### Muleba\_arabiensis\_Delta\_2R\_28335316

##### Muleba\_arabiensis\_Delta\_2R\_28366295

##### Muleba\_arabiensis\_Delta\_2R\_28439507

##### Muleba\_arabiensis\_Delta\_2R\_28616955

---

#### Moshi\_*arabiensis*\_PM

Moshi\_arabiensis\_PM\_2R:24337928

##### Moshi\_arabiensis\_PM\_2R\_24337928

---
