## Supplementary Data S3 for "Copy number variants underlie the major selective sweeps in insecticide resistance genes in *Anopheles arabiensis* from Tanzania": Supplementary_Data_S3.html

window\_Fst\_haplotype\_summary


### F_ST_ windows of interest

For each sample set, we first provide a summary plot of
F_ST_ bewteen resistant and susceptible samples across the
genome, with F_ST_ shown in red and the results of the 200
randomisations shown behind in grey. Windows identified as peaks are
highlighted by points, colour-coded by whether they are significantly
higher than expected based on the simulations (green) or not (purple).
For each significant window (green points), we then provide four
additional figures. The first shows the significant (*P* <
0.01) SNPs found in the region of that window, and their -log10(Pvalue)
of association with phenotype. Red points indicate non-synonymous SNPs,
blue points indicate all other SNPs. Point shape indicates whether the
mutant allele at that SNP is associated with increased (circle) or
decreased (triangle) resistance. Dark points in the centre of the plot
show SNPs within the significant window, light points on the sides show
SNPs in region 10,000 bp either side of the window. The second plot
shows the hierarchical cluster of haplotypes in the window, with
brackets indicating large haplotype clusters. The third plot shows, each
haplotype cluster, indicating the number of haplotypes comprising the
cluster (n), the *P* value of association of the cluster with
resistance phenotype, and the SNPs that differentiate that cluster from
other haplotypes in the region, colour-coded by type (red =
non-synonymous, yellow = synonymous, blue = non-coding). The final
figure shows the contingency tables of phenotype association for each
haplotype cluster.

Legend  
Moshi\_*arabiensis*\_Delta  
Muleba\_*arabiensis*\_Delta  
Moshi\_*arabiensis*\_PM

---

#### Plot legend

---

#### Moshi\_*arabiensis*\_Delta

Moshi\_arabiensis\_Delta\_2R:30957775-30988675

##### Moshi\_arabiensis\_Delta\_2R\_30957775-30988675

###### Contingency table for cluster\_1:

| phenotype | wt | het | hom |
| --- | --- | --- | --- |
| alive | 51 | 27 | 2 |
| dead | 39 | 11 | 1 |

###### Contingency table for cluster\_2:

| phenotype | wt | het | hom |
| --- | --- | --- | --- |
| alive | 70 | 10 | 0 |
| dead | 34 | 13 | 4 |

---

#### Muleba\_*arabiensis*\_Delta

Muleba\_arabiensis\_Delta\_2R:27638426-27662872  
Muleba\_arabiensis\_Delta\_2R:27685539-27705619  
Muleba\_arabiensis\_Delta\_2R:27816983-27842983  
Muleba\_arabiensis\_Delta\_2R:27842983-27881640  
Muleba\_arabiensis\_Delta\_2R:27881639-27914408  
Muleba\_arabiensis\_Delta\_2R:27914407-27940942  
Muleba\_arabiensis\_Delta\_2R:28053952-28093398  
Muleba\_arabiensis\_Delta\_2R:28093397-28143756  
Muleba\_arabiensis\_Delta\_2R:28143756-28184032  
Muleba\_arabiensis\_Delta\_2R:28184031-28228003  
Muleba\_arabiensis\_Delta\_2R:28228002-28259252  
Muleba\_arabiensis\_Delta\_2R:28259251-28286905  
Muleba\_arabiensis\_Delta\_2R:28286905-28322651  
Muleba\_arabiensis\_Delta\_2R:28322651-28359439  
Muleba\_arabiensis\_Delta\_2R:28861167-28883982  
Muleba\_arabiensis\_Delta\_2R:56192837-56228671  
Muleba\_arabiensis\_Delta\_2R:56720537-56789287

##### Muleba\_arabiensis\_Delta\_2R\_27638426-27662872

###### Contingency table for cluster\_1:

| phenotype | wt | het | hom |
| --- | --- | --- | --- |
| alive | 28 | 41 | 12 |
| dead | 49 | 25 | 7 |

###### Contingency table for cluster\_2:

| phenotype | wt | het | hom |
| --- | --- | --- | --- |
| alive | 60 | 20 | 1 |
| dead | 52 | 26 | 3 |

##### Muleba\_arabiensis\_Delta\_2R\_27685539-27705619

###### Contingency table for cluster\_1:

| phenotype | wt | het | hom |
| --- | --- | --- | --- |
| alive | 31 | 38 | 12 |
| dead | 48 | 27 | 6 |

###### Contingency table for cluster\_2:

| phenotype | wt | het | hom |
| --- | --- | --- | --- |
| alive | 62 | 17 | 2 |
| dead | 50 | 28 | 3 |

##### Muleba\_arabiensis\_Delta\_2R\_27816983-27842983

###### Contingency table for cluster\_1:

| phenotype | wt | het | hom |
| --- | --- | --- | --- |
| alive | 28 | 38 | 15 |
| dead | 44 | 30 | 7 |

###### Contingency table for cluster\_2:

| phenotype | wt | het | hom |
| --- | --- | --- | --- |
| alive | 58 | 22 | 1 |
| dead | 48 | 28 | 5 |

###### Contingency table for cluster\_3:

| phenotype | wt | het | hom |
| --- | --- | --- | --- |
| alive | 72 | 9 | 0 |
| dead | 67 | 14 | 0 |

##### Muleba\_arabiensis\_Delta\_2R\_27842983-27881640

###### Contingency table for cluster\_1:

| phenotype | wt | het | hom |
| --- | --- | --- | --- |
| alive | 28 | 37 | 16 |
| dead | 45 | 30 | 6 |

###### Contingency table for cluster\_2:

| phenotype | wt | het | hom |
| --- | --- | --- | --- |
| alive | 58 | 21 | 2 |
| dead | 48 | 28 | 5 |

##### Muleba\_arabiensis\_Delta\_2R\_27881639-27914408

###### Contingency table for cluster\_1:

| phenotype | wt | het | hom |
| --- | --- | --- | --- |
| alive | 53 | 28 | 0 |
| dead | 60 | 21 | 0 |

###### Contingency table for cluster\_2:

| phenotype | wt | het | hom |
| --- | --- | --- | --- |
| alive | 46 | 35 | 0 |
| dead | 71 | 10 | 0 |

###### Contingency table for cluster\_3:

| phenotype | wt | het | hom |
| --- | --- | --- | --- |
| alive | 68 | 13 | 0 |
| dead | 61 | 20 | 0 |

###### Contingency table for cluster\_4:

| phenotype | wt | het | hom |
| --- | --- | --- | --- |
| alive | 73 | 8 | 0 |
| dead | 67 | 14 | 0 |

##### Muleba\_arabiensis\_Delta\_2R\_27914407-27940942

###### Contingency table for cluster\_1:

| phenotype | wt | het | hom |
| --- | --- | --- | --- |
| alive | 26 | 38 | 17 |
| dead | 44 | 31 | 6 |

###### Contingency table for cluster\_2:

| phenotype | wt | het | hom |
| --- | --- | --- | --- |
| alive | 59 | 20 | 2 |
| dead | 47 | 29 | 5 |

##### Muleba\_arabiensis\_Delta\_2R\_28053952-28093398

###### Contingency table for cluster\_1:

| phenotype | wt | het | hom |
| --- | --- | --- | --- |
| alive | 22 | 41 | 18 |
| dead | 42 | 31 | 8 |

###### Contingency table for cluster\_2:

| phenotype | wt | het | hom |
| --- | --- | --- | --- |
| alive | 57 | 21 | 3 |
| dead | 39 | 34 | 8 |

###### Contingency table for cluster\_3:

| phenotype | wt | het | hom |
| --- | --- | --- | --- |
| alive | 69 | 11 | 1 |
| dead | 70 | 11 | 0 |

###### Contingency table for cluster\_4:

| phenotype | wt | het | hom |
| --- | --- | --- | --- |
| alive | 72 | 9 | 0 |
| dead | 66 | 15 | 0 |

##### Muleba\_arabiensis\_Delta\_2R\_28093397-28143756

###### Contingency table for cluster\_1:

| phenotype | wt | het | hom |
| --- | --- | --- | --- |
| alive | 22 | 39 | 20 |
| dead | 41 | 32 | 8 |

###### Contingency table for cluster\_2:

| phenotype | wt | het | hom |
| --- | --- | --- | --- |
| alive | 57 | 20 | 4 |
| dead | 40 | 34 | 7 |

###### Contingency table for cluster\_3:

| phenotype | wt | het | hom |
| --- | --- | --- | --- |
| alive | 72 | 9 | 0 |
| dead | 65 | 16 | 0 |

###### Contingency table for cluster\_4:

| phenotype | wt | het | hom |
| --- | --- | --- | --- |
| alive | 71 | 10 | 0 |
| dead | 70 | 11 | 0 |

##### Muleba\_arabiensis\_Delta\_2R\_28143756-28184032

###### Contingency table for cluster\_1:

| phenotype | wt | het | hom |
| --- | --- | --- | --- |
| alive | 19 | 39 | 23 |
| dead | 38 | 35 | 8 |

###### Contingency table for cluster\_2:

| phenotype | wt | het | hom |
| --- | --- | --- | --- |
| alive | 57 | 20 | 4 |
| dead | 42 | 32 | 7 |

###### Contingency table for cluster\_3:

| phenotype | wt | het | hom |
| --- | --- | --- | --- |
| alive | 71 | 10 | 0 |
| dead | 64 | 16 | 1 |

##### Muleba\_arabiensis\_Delta\_2R\_28184031-28228003

###### Contingency table for cluster\_1:

| phenotype | wt | het | hom |
| --- | --- | --- | --- |
| alive | 18 | 39 | 24 |
| dead | 36 | 37 | 8 |

###### Contingency table for cluster\_2:

| phenotype | wt | het | hom |
| --- | --- | --- | --- |
| alive | 55 | 21 | 5 |
| dead | 41 | 32 | 8 |

###### Contingency table for cluster\_3:

| phenotype | wt | het | hom |
| --- | --- | --- | --- |
| alive | 71 | 10 | 0 |
| dead | 64 | 17 | 0 |

###### Contingency table for cluster\_4:

| phenotype | wt | het | hom |
| --- | --- | --- | --- |
| alive | 70 | 11 | 0 |
| dead | 68 | 13 | 0 |

##### Muleba\_arabiensis\_Delta\_2R\_28228002-28259252

###### Contingency table for cluster\_1:

| phenotype | wt | het | hom |
| --- | --- | --- | --- |
| alive | 15 | 42 | 24 |
| dead | 33 | 39 | 9 |

###### Contingency table for cluster\_2:

| phenotype | wt | het | hom |
| --- | --- | --- | --- |
| alive | 56 | 20 | 5 |
| dead | 44 | 29 | 8 |

###### Contingency table for cluster\_3:

| phenotype | wt | het | hom |
| --- | --- | --- | --- |
| alive | 72 | 9 | 0 |
| dead | 66 | 15 | 0 |

##### Muleba\_arabiensis\_Delta\_2R\_28259251-28286905

###### Contingency table for cluster\_1:

| phenotype | wt | het | hom |
| --- | --- | --- | --- |
| alive | 15 | 42 | 24 |
| dead | 30 | 40 | 11 |

###### Contingency table for cluster\_2:

| phenotype | wt | het | hom |
| --- | --- | --- | --- |
| alive | 54 | 22 | 5 |
| dead | 42 | 31 | 8 |

###### Contingency table for cluster\_3:

| phenotype | wt | het | hom |
| --- | --- | --- | --- |
| alive | 72 | 9 | 0 |
| dead | 67 | 14 | 0 |

##### Muleba\_arabiensis\_Delta\_2R\_28286905-28322651

###### Contingency table for cluster\_1:

| phenotype | wt | het | hom |
| --- | --- | --- | --- |
| alive | 12 | 44 | 25 |
| dead | 27 | 42 | 12 |

###### Contingency table for cluster\_2:

| phenotype | wt | het | hom |
| --- | --- | --- | --- |
| alive | 52 | 24 | 5 |
| dead | 40 | 33 | 8 |

###### Contingency table for cluster\_3:

| phenotype | wt | het | hom |
| --- | --- | --- | --- |
| alive | 73 | 8 | 0 |
| dead | 67 | 14 | 0 |

##### Muleba\_arabiensis\_Delta\_2R\_28322651-28359439

###### Contingency table for cluster\_1:

| phenotype | wt | het | hom |
| --- | --- | --- | --- |
| alive | 12 | 45 | 24 |
| dead | 27 | 41 | 13 |

###### Contingency table for cluster\_2:

| phenotype | wt | het | hom |
| --- | --- | --- | --- |
| alive | 52 | 24 | 5 |
| dead | 42 | 32 | 7 |

###### Contingency table for cluster\_3:

| phenotype | wt | het | hom |
| --- | --- | --- | --- |
| alive | 72 | 9 | 0 |
| dead | 68 | 13 | 0 |

##### Muleba\_arabiensis\_Delta\_2R\_28861167-28883982

###### Contingency table for cluster\_1:

| phenotype | wt | het | hom |
| --- | --- | --- | --- |
| alive | 35 | 42 | 4 |
| dead | 33 | 38 | 10 |

###### Contingency table for cluster\_2:

| phenotype | wt | het | hom |
| --- | --- | --- | --- |
| alive | 45 | 32 | 4 |
| dead | 40 | 35 | 6 |

##### Muleba\_arabiensis\_Delta\_2R\_56192837-56228671

###### Contingency table for cluster\_1:

| phenotype | wt | het | hom |
| --- | --- | --- | --- |
| alive | 42 | 36 | 3 |
| dead | 56 | 21 | 4 |

###### Contingency table for cluster\_2:

| phenotype | wt | het | hom |
| --- | --- | --- | --- |
| alive | 67 | 11 | 3 |
| dead | 62 | 17 | 2 |

###### Contingency table for cluster\_3:

| phenotype | wt | het | hom |
| --- | --- | --- | --- |
| alive | 69 | 12 | 0 |
| dead | 61 | 19 | 1 |

###### Contingency table for cluster\_4:

| phenotype | wt | het | hom |
| --- | --- | --- | --- |
| alive | 71 | 9 | 1 |
| dead | 61 | 20 | 0 |

###### Contingency table for cluster\_5:

| phenotype | wt | het | hom |
| --- | --- | --- | --- |
| alive | 72 | 8 | 1 |
| dead | 71 | 8 | 2 |

###### Contingency table for cluster\_6:

| phenotype | wt | het | hom |
| --- | --- | --- | --- |
| alive | 66 | 15 | 0 |
| dead | 75 | 6 | 0 |

##### Muleba\_arabiensis\_Delta\_2R\_56720537-56789287

###### Contingency table for cluster\_1:

| phenotype | wt | het | hom |
| --- | --- | --- | --- |
| alive | 67 | 11 | 3 |
| dead | 62 | 16 | 3 |

###### Contingency table for cluster\_2:

| phenotype | wt | het | hom |
| --- | --- | --- | --- |
| alive | 70 | 11 | 0 |
| dead | 59 | 20 | 2 |

###### Contingency table for cluster\_3:

| phenotype | wt | het | hom |
| --- | --- | --- | --- |
| alive | 71 | 9 | 1 |
| dead | 60 | 21 | 0 |

###### Contingency table for cluster\_4:

| phenotype | wt | het | hom |
| --- | --- | --- | --- |
| alive | 58 | 22 | 1 |
| dead | 73 | 8 | 0 |

###### Contingency table for cluster\_5:

| phenotype | wt | het | hom |
| --- | --- | --- | --- |
| alive | 61 | 20 | 0 |
| dead | 70 | 10 | 1 |

###### Contingency table for cluster\_6:

| phenotype | wt | het | hom |
| --- | --- | --- | --- |
| alive | 69 | 11 | 1 |
| dead | 72 | 6 | 3 |

###### Contingency table for cluster\_7:

| phenotype | wt | het | hom |
| --- | --- | --- | --- |
| alive | 69 | 12 | 0 |
| dead | 72 | 9 | 0 |

---

#### Moshi\_*arabiensis*\_PM

Moshi\_arabiensis\_PM\_3R:46657236-46697631

##### Moshi\_arabiensis\_PM\_3R\_46657236-46697631

###### Contingency table for cluster\_1:

| phenotype | wt | het | hom |
| --- | --- | --- | --- |
| alive | 52 | 26 | 4 |
| dead | 54 | 15 | 0 |

###### Contingency table for cluster\_2:

| phenotype | wt | het | hom |
| --- | --- | --- | --- |
| alive | 69 | 12 | 1 |
| dead | 57 | 12 | 0 |

###### Contingency table for cluster\_3:

| phenotype | wt | het | hom |
| --- | --- | --- | --- |
| alive | 71 | 11 | 0 |
| dead | 61 | 5 | 3 |

---
